# Reconstructing the history of polygenic scores using coalescent trees

**DOI:** 10.1101/389221

**Authors:** Michael D. Edge, Graham Coop

## Abstract

Genome-wide association studies (GWAS) have revealed that many traits are highly polygenic, in that their within-population variance is governed in part by small-effect variants at many genetic loci. Standard population-genetic methods for inferring evolutionary history are ill-suited for polygenic traits—when there are many variants of small effect, signatures of natural selection are spread across the genome and subtle at any one locus. In the last several years, several methods have emerged for detecting the action of natural selection on polygenic scores, sums of genotypes weighted by GWAS effect sizes. However, most existing methods do not reveal the timing or strength of selection. Here, we present a set of methods for estimating the historical time course of a population-mean polygenic score using local coalescent trees at GWAS loci. These time courses are estimated by using coalescent theory to relate the branch lengths of trees to allele-frequency change. The resulting time course can be tested for evidence of natural selection. We present theory and simulations supporting our procedures, as well as estimated time courses of polygenic scores for human height. Because of its grounding in coalescent theory, the framework presented here can be extended to a variety of demographic scenarios, and its usefulness will increase as both GWAS and ancestral recombination graph (ARG) inference continue to progress.

## 2 Introduction

Some of the most compelling examples of phenotypic evolution come from time courses that reveal the pace of evolution, either through observations across generations (Cook et al., 1986; Grant and Grant, 2002) or through changes in the fossil record (Gingerich, 1983; MacFadden, 2005; Bell et al., 2006). For many traits and species, it can be difficult to ascertain whether these changes reflect genetic change. For example, we have fairly detailed knowledge of human height through time, but some changes in height are likely driven by environmental and dietary changes (Stulp and Barrett, 2016). Thanks to ancient DNA, we can now sometimes obtain a partial picture of long-term genetic changes involving relatively simple traits like pigmentation (Ludwig et al., 2009) or more complex traits (Mathieson et al., 2015). However, we are usually not fortunate enough to have access to genotype data from across time, and even when ancient DNA are available, the resulting time courses will necessarily be incomplete.

One alternative to direct measurement of phenotypes through time is to reconstruct the history of a phenotype using contemporary genetic data. Positive selection on simple genetic traits drives large allele-frequency changes at the causal loci and linked neutral alleles (Smith and Haigh, 1974). There are many procedures for detecting this kind of selection on individual alleles and for dating and modeling their spread through populations (Tajima, 1989; Fay and Wu, 2000; Sabeti et al., 2002; Voight et al., 2006; Ronen et al., 2013; Garud et al., 2015; Crawford et al., 2017).

One obstacle to understanding the evolutionary basis of phenotypes is the polygenic architecture of many traits. Complex traits—traits affected by many genetic loci and by environmental variation—are ill-suited to study by single-locus methods. In recent years, genome-wide association studies (GWAS) have made it possible to aggregate subtle evolutionary signals that are distributed across the many genetic loci that influence a trait of interest (Turchin et al., 2012; Berg and Coop, 2014; Robinson et al., 2015; Field et al., 2016; Racimo et al., 2018; Uricchio et al., 2018). For example, Field et al. developed the singleton density score (or SDS) to infer recent selection on a variety of traits among the ancestors of people in the United Kingdom (Field et al., 2016). (As discussed below, some empirical findings of these studies have not replicated using effect sizes estimates from less-structured GWAS samples, raising the possibility that the selection tests are sensitive to residual population stratification (Berg et al., 2018; Sohail et al., 2018). Nonetheless, given correct effect size estimates, these methods are useful.)

Field et al. relied on the fact that selection distorts the gene genealogies, or coalescent trees, at genetic loci under selection. In particular, loci under positive selection will have increased in frequency in the recent past, leading to relatively faster coalescence of lineages than if the allele frequency had been constant. The principle on which the singleton density score relies is quite general—selection, even when its effect is spread over many loci, leaves systematic signals in coalescent trees at loci underlying trait variation.

The ancestral recombination graph (ARG, Griffiths and Marjoram (1997)) collects coalescent trees at loci along a recombining sequence, encoding information about allele-frequency changes at each site, as well as recombination events between sites. Recently, computational methods for inferring ARGs have advanced considerably (Rasmussen et al., 2014; Mirzaei and Wu, 2016), allowing a range of applications (Palacios et al., 2015).

In this work, we consider ways in which ARGs—and in particular, the coalescent trees of sites associated with a phenotype—might be used to reconstruct the history of the phenotype they influence. The ARG-based approaches we consider are motivated by polygenic traits, and the population-mean level of a polygenic score—a prediction of phenotype from an individual’s genotype—is the target of estimation. We present methods for estimating the time course of a population-mean polygenic score through the past, as well as a test for assessing whether an estimated time course is consistent with neutral evolution alone.

We begin by describing estimators and a hypothesis test for phenotypic time courses based on previous theory. Next, we apply these procedures to simulated data, using both true and reconstructed ARGs. Finally, we apply our methods to some human height in the GBR (Great Britain) subset of the 1000 Genomes panel (1000 Genomes Project Consortium, 2012), using ARGs inferred by RENT+ (Mirzaei and Wu, 2016).

## 3 Theory

**Background and motivation**. The ARG expresses the shared genealogical history of a sample of individuals at a set of genetic loci, accounting for correlations among neighboring loci that arise because of linkage. The ARG contains a coalescent tree for every site in the genome—these trees for specific sites are *marginal trees*. Our methods make use of the marginal trees at a set of sites that are associated with a phenotype. In particular, we concentrate on the information revealed by the number of coalescent lineages that remain (i.e., that have not yet coalesced) at a time *t* in the past. The number of lineages at a given time in the past is described by a stochastic process called the ancestral process (see e.g. Tavaré, 1984).

The intuition behind the methods we present here is shown in Figure 1. If an allele has been selected upward in frequency in the recent past, then the number of chromosomes carrying the allele will likely have increased. Looking backward in time, the number of carriers in the recent past is less than in the present, which forces an excess of coalescence events recently. Similarly, if an allele has been selected downward recently, then there will be tend to be fewer recent coalescent events compared with the neutral expectation. If the trait that has been under selection is polygenic, then the signal at each locus associated with the trait will be smaller, and its strength and direction will depend on the effect size at the locus. In Appendix A, we derive the relationship between the rate of coalescence and selection on the phenotype. We show that phenotypic selection acting to increase the population-mean trait value increases the rate of coalescence for alleles that increase the trait and lowers the coalescence rate for alleles that decrease the trait. In contrast, stabilizing selection acting on a trait for which the population mean is at the fitness optimum does not have a systematic directional effect on the coalescent rates.

**Figure 1:**
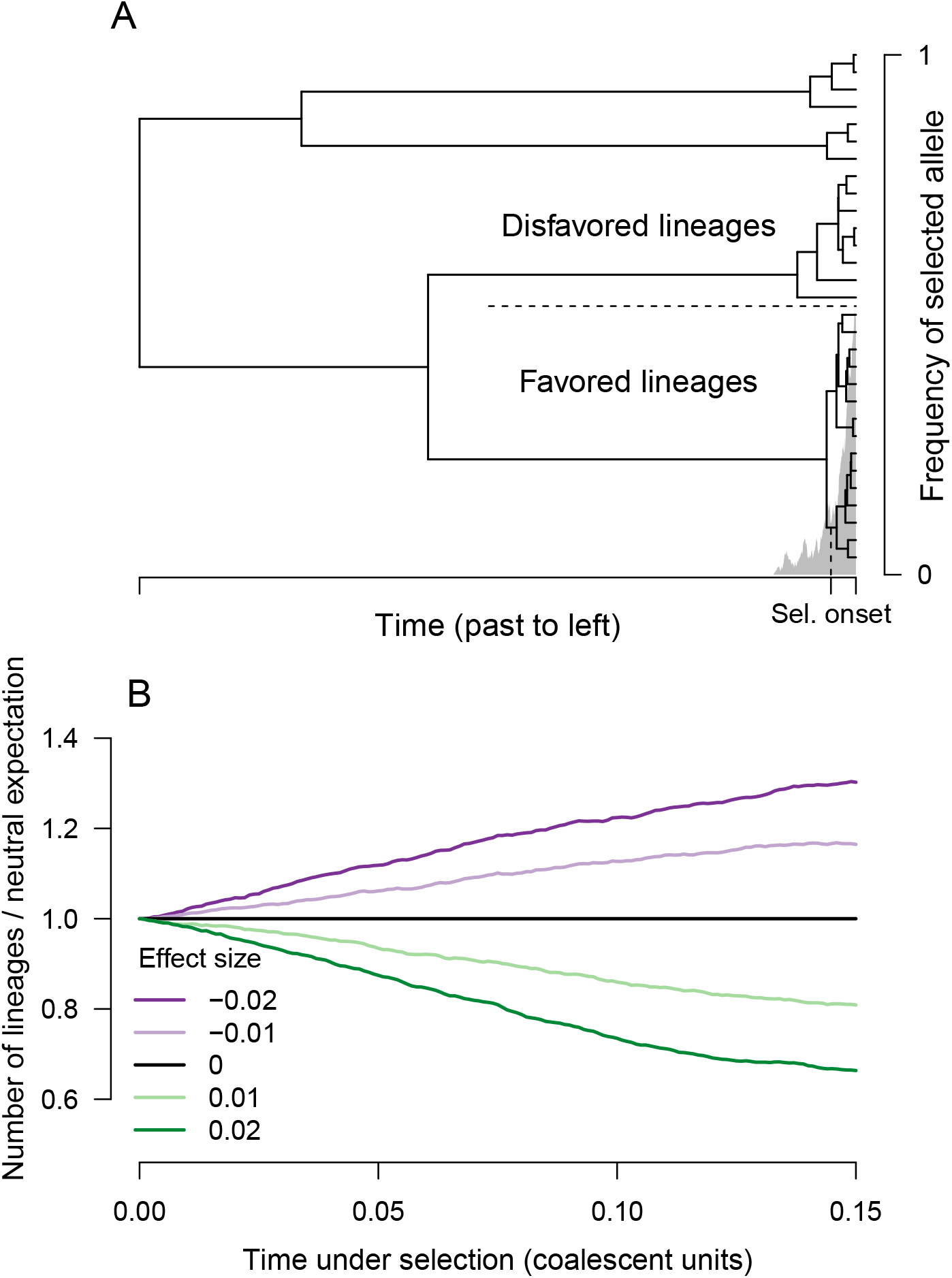
Selection distorts the coalescent tree at a locus. A) The frequency time course for an allele that has been favored by recent selection is shown in gray. Superimposed is a coalescent tree relating a sample of chromosomes; the top half are of the disfavored type, and the bottom half are of the favored type. Because the favored alleles trace to a pool of ancestors that was small before selection, there is an excess of recent coalescence on the subtree of the favored allele. B) If a polygenic trait is under directional selection, then the rate of coalescence depends on effect size. The time under selection is shown on the horizontal axis, and the vertical axis shows the ratio of lineages remaining compared with the neutral expectation. Different lines show different effect sizes in units of trait standard deviation. Results are an average across 1,000 simulations; selection coefficients are determined as if the trait experiences 1% directional truncation selection.

Our target of estimation is the population-average polygenic score for a trait going backward through time. By “polygenic score,” we mean a weighted sum of an individual’s genotypes, where the weights are the additive effect sizes of each allele. In our case, we are interested in the population-average polygenic score, so we take a weighted sum of the allele frequencies,

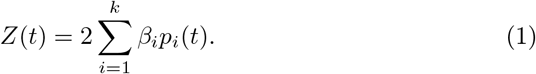

Here, *Z*(*t*) is the population-average polygenic score at time *t* in the past, *p_i_*(*t*) is the population frequency of one of the two alleles at locus *i* at time *t* in the past, and *β_i_* is the additive effect size of the allele whose frequency is *P_i_*(*t*), where the effect sizes have been scaled so that the other allele has an effect size of zero. (In practice, effect sizes will be estimated with error, but in this paper we treat the effect sizes as known.) The 2 arises because of diploidy.

If the *k* loci included in the calculation of *Z*(*t*) include all the causal loci, then changes in *Z*(*t*), the population-average polygenic score, reflect changes in the population-average phenotypic value in the absence of changes in the distribution of environmental effects on the trait, changes in the effect size, epistasis, and gene-by-environment interaction. Even if these strong assumptions are not met, then rapid changes in *Z*(*t*) could provide evidence that natural selection has acted on the trait.

Our strategy for estimating *Z*(*t*) is to estimate the historical allele-frequency time courses, *p_i_*(*t*), for the loci associated with a trait. Given an estimator of the allele-frequency time courses, 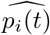, we estimate polygenic scores as

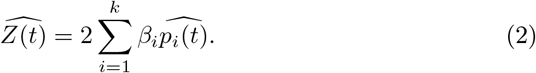

If the loci contributing to the polygenic score are independent, then the variance of the polygenic score estimator is

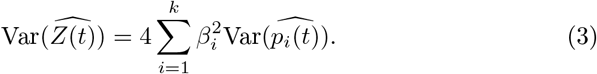

We present three methods for estimating historical allele-frequency time courses. A number of authors have investigated estimating allele-frequency time courses from coalescent genealogies (Slatkin, 2001; Coop and Griffiths, 2004; Chen and Slatkin, 2013) or applying Wright-Fisher diffusion theory to time-series data (Bollback et al., 2008; Schraiber et al., 2016). Our approaches are cruder than some of these but have the advantage of being fast enough to be applicable to thousands of GWAS loci.

### 3.1 Estimating the allele-frequency time course at a single locus

We present several methods for estimating the historical allele-frequency time course at a specific biallelic locus given a coalescent tree at the locus. (Our procedures could be generalized to loci with multiple alleles.) In each case, the goal is to estimate the frequency of an allele of interest (e.g. the effect allele) at locus *i* at time *t* in the past, or *p_i_*(*t*).

#### 3.1.1 Proportion-of-lineages estimator

The simplest way to estimate a historical allele frequency is to treat the lineages ancestral to the sample at time *t* as representative of the population at time *t*. If the locus has evolved neutrally between the present and time *t* in the past, then the lineages ancestral to the sample at time *t* are a random sample—with respect to allelic type— from the population at time *t*. If the lineages ancestral to the sample are a random sample from the population at time *t*, then a reasonable estimator of *p_i_*(*t*) is the proportion of lineages at time *t* that carry the allele of interest,

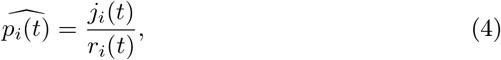

where *r_i_*(*t*) is the total number of lineages ancestral to the present-day sample at locus *i* at time *t*, and *j_i_*(*t*) is the number of lineages at time *t* that carry the allele of interest. Assuming that the mutation distinguishing the alleles has only appeared once in the history of the sample, the lineages that carry the derived allele are those ancestral to contemporary copies of the derived allele, which must coalesce to one lineage before coalescing with the rest of the tree. If the tree for the locus is known, then the branch on which the mutation must have appeared is known, but the exact time of the mutation is not. In practice, we assume that the mutation occurred in the middle of the branch connecting the derived subtree to the rest of the tree. (We make this assumption when implementing all estimators.)

**Figure 2:**
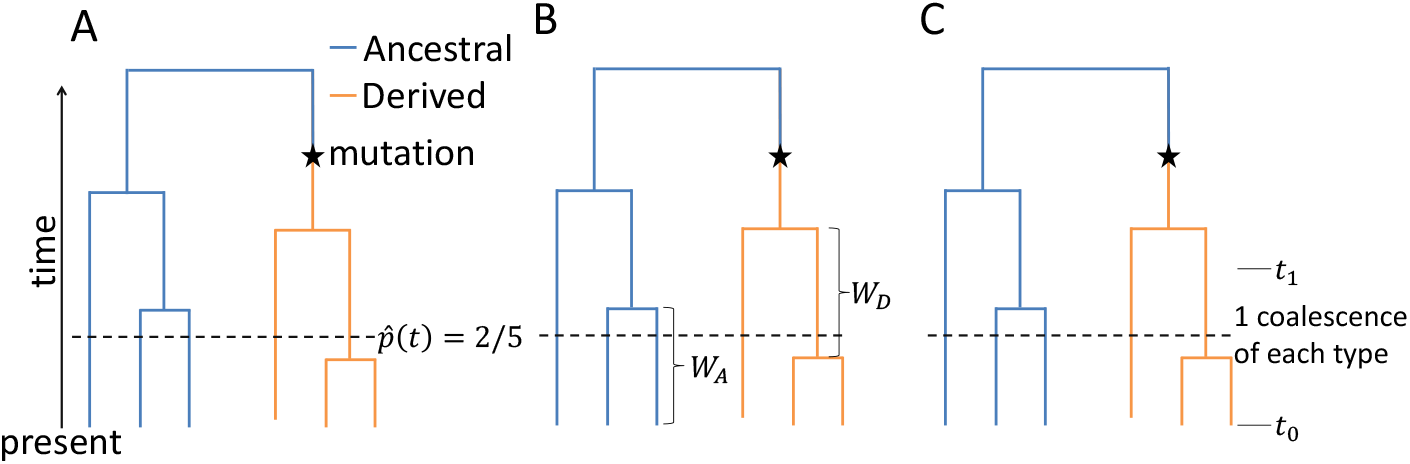
Schematics of three estimators of historical allele frequencies using coalescent trees. In each tree, we show the information used to estimate allele frequency in the population at a time represented by the horizontal dashed line. A) The proportion-of-lineages estimator. The estimate of the derived allele frequency in the population is the derived allele frequency among the ancestors of the sample, or 2/5 in the picture. B) The waiting-time estimator. Relative “sizes” of the subpopulations of ancestral-allele and derived-allele carriers are estimated as a function of intercoalescence times among the ancestors of the present-day ancestral- and derived-allele carriers. C) The lineages-remaining estimator. The relative sizes of the subpopulations of ancestral-allele and derived-allele carriers are estimated by examining the number of coalescence events that occur between pre-specified time points, here called *t*_0_ and *t*_1_.

If the population size at time *t* is large compared with the number of ancestral lineages at time t, then conditional on *r_i_*(*t*), the number of ancestral lineages carrying the allele of interest, *j_i_*(*t*), is distributed approximately as a Binomial(*r_i_*(*t*), *p_i_*(*t*)) random variable. Thus, Eq. 4 is the maximum-likelihood estimator of *p_i_*(*t*), and conditional on *r_i_*(*t*), its sampling variance can be estimated as (dropping the subscript is and parenthetical ts for compactness)

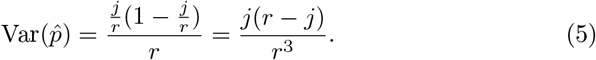

In appendix B, we give a Bayesian interpretation of the proportion-of-lineages estimator that relies on connections between neutral diffusion and the ancestral process (Tavaré, 1984).

If chromosomes carrying the two alleles have differed in fitness between the present and time t, then Eq. 4 will in general be a biased and inconsistent estimator of *p_i_*(*t*). Chromosomes carrying alleles that have been favored by selection will be more likely to leave offspring in the present-day population than will chromosomes carrying unfavored alleles. Favored alleles will thus be overrepresented among the lineages ancestral to the sample compared with their actual frequency at time *t*. However, even if Eq. 4 is an inconsistent estimator of the population allele frequency, it retains the interpretation of being the allele frequency among lineages ancestral to the sample. Thus, when Eq. 2 is applied to allele frequencies estimated by Eq. 4, the result is the mean polygenic score among chromosomes ancestral to the present-day sample at some time in the past.

#### 3.1.2 Waiting-time estimator

The proportion-of-lineages estimator proposed in Eq. 4 works well under neutrality, but under selection, it tends to underestimate the degree of allele-frequency change experienced by selected alleles. One potential solution is to consider the chromosomes in the population as two separate subpopulations (Hudson and Kaplan, 1988)—one for the carriers of each of the two alleles at the locus—and to estimate the sizes of those two populations over time. At locus *i*, denote the sizes of these two subpopulations at time *t* as *N_i_*(*t*) (for the allele of interest) and *M_i_*(*t*) (for the other allele). The frequency of the allele of interest at time *t* in the past, or *p_i_*(*t*), is

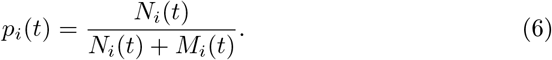

Here, we propose to estimate *N_i_*(*t*) and *M_i_*(*t*) on the basis of properties of the coalescent trees for the two alleles. We then estimate *p_i_*(*t*) by plugging these estimates, 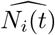 and 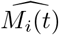, into Eq. 6, giving

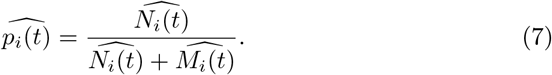

The estimator in Eq. 7 is not unbiased in general, even if the estimates of *N_i_*(*t*) and *M_i_*(*t*) are unbiased, but we give one justification of Eq. 7 via a first-order Taylor-series approximation in Appendix C. Further, separating the problems of estimating the two subpopulations has an advantage—this estimator does not assume, as the proportion-of-lineages estimator does, that the two allelic types have had equal fitness between *t* and the present.

Assuming that the mutation that distinguished the two alleles occurred only once in the history of the sample, the chromosomes carrying the two alleles can be treated as two distinct subsamples between the time of the mutation and the present. Among the ancestors of the allele subsample carrying the allele of interest, coalescent time *τ* accrues according to

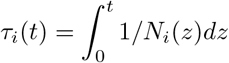

where *t* is measured in generations. It follows that

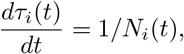

suggesting that *N_i_*(*t*) can be estimated by assessing the rate at which coalescent time accrues. *M_i_*(*t*) can then be estimated analogously, and *p_i_*(*t*) can be estimated by Eq. 7. We present two related approaches to estimating the rate of accrual of coalescent time in each subsample—one approach in which estimates are made with respect to waiting times between coalescent events, and another in which estimates are made with respect to the number of lineages ancestral to the subsample at a specified time in the past. In both approaches, we assume that *N_i_*(*t*) and *M_i_*(*t*) are piecewise constant, but it is possible to modify these estimators under other assumptions about how *N_i_*(*t*) and *M_i_*(*t*) change between timepoints.

The number of coalescence events in a time interval depends on the amount of coalescent time passed. Our approaches amount to inverting this relationship to estimate the amount of coalescent time passed on each lineage (and thus their relative population sizes) in a method-of-moments framework. In this subsection, we assess *N_i_*(*t*) and *M_i_*(*t*) according to the time passed between fixed numbers of coalescent events.

Suppose that *N_i_*(*t*) = *N*, assumed to be constant from a timepoint of interest (defined here to be *t* = 0) until *ℓ* coalescences have occured within the subsample, which at *t* = 0 consists of *n_i_* lineages. Define *Y_k_* as the waiting time to the *k*th coalescence, starting from the *k* – 1th coalesence event (or from *t* = 0 if *k* = 1). Define the total waiting time from *t* = 0 to the *ℓ*-th coalescence as 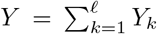. Then in units of generations, each *Y_k_* is an independent, exponentially distributed random variable with rate (*n_i_* − *k* + 1)(*n_i_* − *k*)/(2*N*). Thus,

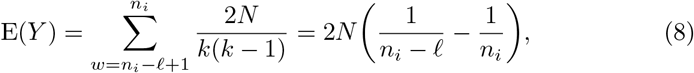

and

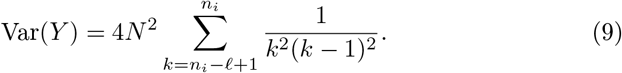

For large *n_i_* and intermediate values of *ℓ*, *Y* is approximately normally distributed with expectation and variance given by Eqs. 8 and 9 (Chen and Chen, 2013). One estimator of *N_i_*(*t*) = *N*, which is both a method-of-moments estimator and the maximum-likelihood estimator under the asymptotically normal distribution of *Y*, is

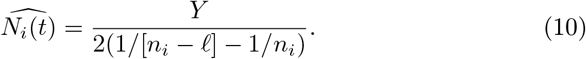

With *ℓ* =1, this is the “skyline” estimator of Pybus et al. (2000), and with *ℓ* ≥ 1, it is the generalized skyline estimator of Strimmer and Pybus (2001). It is unbiased under the assumption of constant *N_i_*(*t*) between coalescence events. In our implementation, after the tree has coalesced down to one lineage, we assume that *N_i_*(*t*) remains constant into the past if its allele is ancestral and that *N_i_*(*t*) remains constant before dropping to zero in the middle of the branch on which the mutation arose if its allele is derived. (In principle, derived vs. ancestral status may be determined by the tree topology or “forced” based on prior knowledge if the tree topology may be in error. We use the tree topology.)

One may estimate the frequency of the a allele at time *t* by

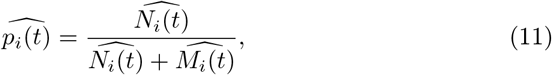

where *M_i_*(*t*) is estimated analogously to *N_i_*(*t*). In principle, the estimates of *N_i_*(*t*) and *M_i_*(*t*) may be based on waiting times for different numbers of coalescences (*ℓ*), and they will likely be constant for different intervals of time. By a first-order Taylor approximation argument described in Appendix C,

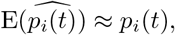

but in practice, 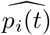 can be substantially biased. Its approximate variance is given in Appendix C.

One important decision in using this estimator is the choice of *ℓ*—i.e., the number of coalescence events to wait for—for each allele. Small values of *ℓ* lead to variable estimates. On the other hand, the estimator assumes that the size of the subpopulation is constant until *ℓ* coalescent events have occurred, which may be increasingly unrealistic for large *ℓ*. In this paper, we use *ℓ* = 1 and defer investigation of different choices of *ℓ* for future work.

#### 3.1.3 Lineages-remaining estimator

The next estimator, which we term the “lineages-remaining” estimator, operates on a principle similar to the waiting-time estimator. The coalescent time passed on each allele’s background is evaluated by looking at the local rate of coalescence, and the relative numbers of carriers of each allele are estimated to form an allele-frequency estimate. The difference is that whereas the waiting-time estimator estimates “population sizes” for each allele between coalescent events, the lineages-remaining estimator estimates the “population sizes” between pre-specified times by comparing the number of lineages of each type that remain (i.e. have not coalesced) at the more ancient end of a time interval with the number present at the more recent end of the interval.

Suppose that *N_i_*(*t*) = *N* during an interval of Δ*t* generations. At the end of the interval closer to the present, there are 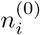 lineages, and at the end of the interval further into the past, there are 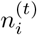 lineages. The expected number of lineages remaining at the end of the interval can be approximated as

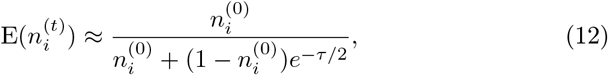

where *τ* is the amount of coalescent-scaled time elapsed during the interval (Griffiths, 1984; Maruvka et al., 2011; Chen and Chen, 2013; Jewett and Rosenberg, 2014). Here, *N_i_*(*t*) is a haploid population size, so *τ* = Δ*t*/*N_i_*(*t*). For large 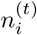 and intermediate lengths of time, is approximately normally distributed (Griffiths, 1984; Chen and Chen, 2013). An estimator for *N_i_*(*t*) = *N*, which is both a method-of-moments estimator (one based on the approximate value of 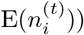 and the maximum-likelihood estimator under the limiting normal distribution, is

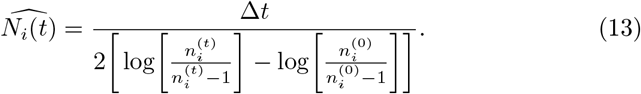

As the tree coalesces to few lineages, there are several edge cases in which the estimator is undefined. Our methods for handling these cases are discussed in Appendix C.

It may be impractical to estimate *N_i_*(*t*) because Δ*t* in generations may be unknown. However, Δt cancels in the estimator of the allele frequency *p_i_*(*t*),

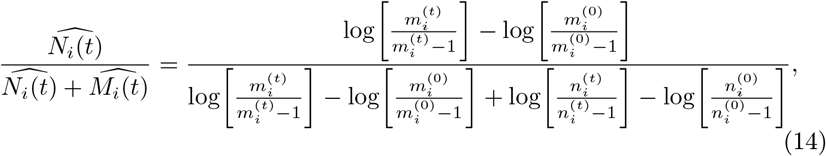

where *M_i_*(*t*) is the number of carriers of the reference allele in the population at locus *i* at time *t*, and *m_i_*(0) and *m_i_*(*t*) are the numbers of lineages carrying the reference allele at the ends of the time interval closer to and more distant from the present, respectively. Approximate expressions for the variance of the estimator in Eq. 14 are in Appendix C.

In practice, the estimator in Eq. 14 will be evaluated at a set of times. Here, we evaluate the estimator every .001 coalescent units, starting at the present and extending back 4 coalescent units. The grid of times at which changes in the number of lineages of each type are considered will influence the estimate. Finer grids will lead to estimates that are more variable but less biased by the assumption of constant *N_i_*(*t*) and *M_i_*(*t*) between time points.

### 3.2 Testing time courses for selection

Once a polygenic score time course has been constructed, it can be tested for selection. Whereas the relevance for trait evolution of the estimators proposed in the previous sections depends on the proportion of trait variance accounted for by the polygenic score, polygenic-score time courses can be tested for selection even if they account for a small proportion of the trait variance. Further, although we apply the test in this section to estimated polygenic score time courses, it is also applicable to measured time-series data on allele frequencies when available.

We propose a test for selection that amounts to a modification of the *Q_X_* framework of Berg and Coop (2014), an analogue of *Q_ST_-F_ST_* tests for phenotypic-selection (Whitlock, 2008). Berg and Coop proposed *Q_X_* to test for overdispersion of polygenic scores among population samples, relative to neutral expectations. Here, we check for overdispersion among a set of timepoints along one population branch. We denote our time-based modification of *Q_X_* as *T_X_*.

Suppose that for each time *t_j_* in a sequence of times, we observe a population-level polygenic score, 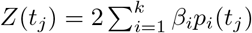, as in Eq. 1. Assume that changes across time at distinct loci are independent. Following the *Q_X_* framework, we model polygenic scores using a multivariate normal distribution. Specifically, we posit that at each locus, over short time scales, allele-frequency changes between timepoints follow a Normal(0, *fp_i_*(1 − *p_i_*)) distribution (Cavalli-Sforza et al., 1964; Nicholson et al., 2002). (Approximate normality fails over longer time scales, in part because allele frequencies are bounded by 0 and 1.) Here, *f* is the coalescent time that has passed between timepoints, and *p_i_* is the allele frequency at locus *i* at one end of the interval. (In practice, we choose *p_i_* to be the allele frequency at the end of the interval closer to the present.) The parameter *f* is constant across loci.

If allele-frequency changes at each locus are independent and normally distributed, then changes between timepoints in the polygenic scores are Normal with expectation 0 and variance 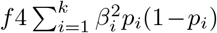. The variance of polygenic score changes can also be written as *f*2*V_A_*, where 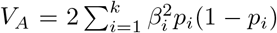 is the additive genetic variance of the trait.

Imagine we have a time course of polygenic scores *Z*(*t*_0_), *Z*(*t*_1_), *Z*(*t*_2_), …, *Z*(*t_w_*), with *t*_0_ < *t*_1_ < … < *t_w_*. Under neutrality, for each timepoint *j* ∈ 1, …, *w*, the statistic

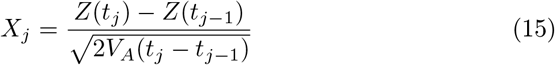

has a Normal(0, 1) distribution. (*V_A_* can be recomputed for each value of *j* using allele frequencies at *t*_*j*−1_. In practice, we use the same *V_A_*—the one computed from allele-frequency estimates closest to the present—for each time interval.) Moreover, under neutrality, allele-frequency changes in distinct time intervals are independent, so values of *X_j_* are independent for distinct *j*. Thus, the sum across timepoints

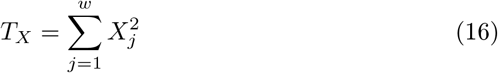

has a *χ*^2^(*w*) distribution. (For details on the equivalence of the *T_X_* statistic in Eq. 16 to the *Q_X_* statistic of Berg and Coop (2014) in this scenario, see Appendix D.) In contrast, under directional selection, changes in allele frequency across time depend on the effect size of the locus, leading to large changes in the polygenic score and *T_X_* values larger than predicted by the *χ*^2^(*w*) distribution. This test is an analogue of time-course tests for phenotypic selection based on neutral Brownian motion (Lande, 1976; Turelli et al., 1988), but with the advantage that we know *V_A_*.

*T_X_* can be compared with a *χ*^2^(*w*) distribution to test for significance, or it can be compared with a distribution obtained by permuting either effect sizes or their signs. If the *χ*^2^(*w*) distribution is used, then coalescent times elapsed between polygenic scores can be estimated by assessing estimated allele-frequency changes between time points, either at putatively neutral loci or trait-associated loci. Using estimated times produces type I error rates closer to the nominal value because the estimators we use tend to change at a systematically different rate than the actual allele frequencies— the proportion-of-lineages estimator changes more slowly than population allele frequencies do, and the other estimators change more quickly than population allele frequencies. In practice, we use the sample variance of 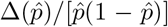 as an estimate of coalescent time passed between timepoints, where 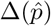 is an estimated allele-frequency change at a variable locus, and 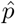 is the estimated allele frequency at the more recent end of the time interval. Assessing elapsed time on the basis of trait-associated loci may lead to a power decrement, but it will not be large— most of the *T_X_* signal comes from coordination of small shifts across loci and not from larger-than-expected allele-frequency changes (Berg and Coop, 2014).

## 4 Simulation results

We examined the performance of our methods in coalescent simulations. In particular, we simulated coalescent trees for unlinked loci associated with a phenotype. The simulated loci evolve neutrally or under directional selection. In particular, if an allele at locus *i* has effect size *β_i_* on a trait, and the trait experiences a selection gradient *α*(*t*) at time *t*, then the selection coefficient on the allele—representing the fitness of the heterozygote minus the fitness of the ancestral homozygote—is *s*(*t*) = *α*(*t*)*β_i_* (Charlesworth and Charlesworth, 2010, Eq. B3.7.7). The coalescent simulations were run in mssel (Berg and Coop, 2015), a version of ms (Hudson, 2002) that takes allele-frequency time courses that may be produced by selection, and assumed a constant population size. Effect sizes for the derived allele are drawn from a normal distribution centered at zero. For details on the simulations, see Appendix E.

We consider the performance of the methods when a) the true trees are provided as input, and when b) the trees must be reconstructed from sequence data. We use the software RENT+ (Mirzaei and Wu, 2016) for tree reconstruction.

Before considering systematic results over many simulations, we present estimated time courses for one representative simulation (Figure 3). In the simulation shown, the trait was under selection upward in the past but has evolved neutrally recently.

**Figure 3:**
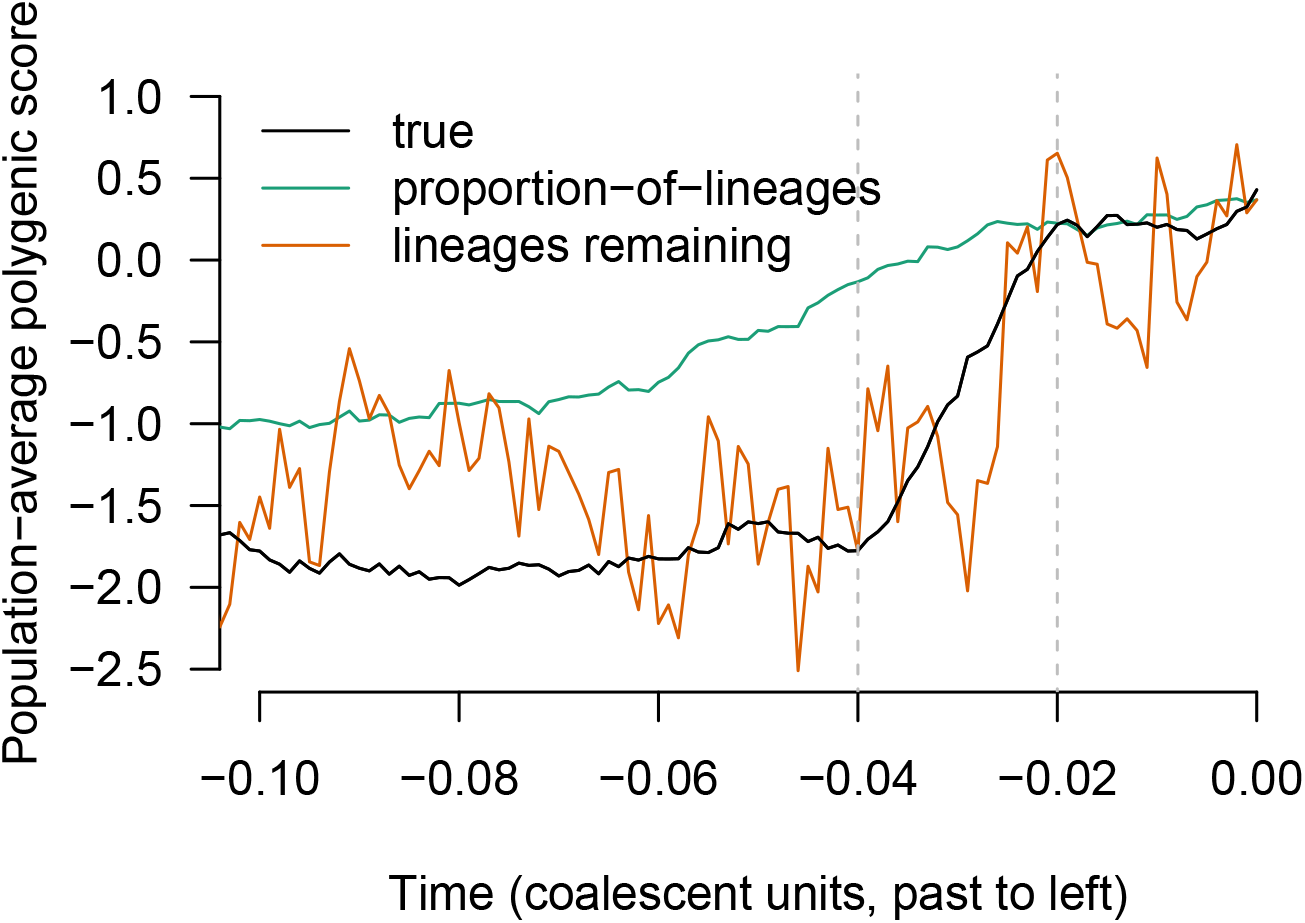
The behavior of two of the proposed estimators—the proportion-of-lineages estimator and the lineages-remaining estimator—in a single simulation. The true time course of the population-average polygenic score is shown in black. The trait is under selection upward from .04 to .02 coalescent units in the past, leading to a shift of approximately two standard deviations in present-day units. Between the present and the most recent period of selection, the proportion-of-lineages estimator performs well. During the period of selection (and before), the ancestors of the sample are no longer representative of the ancestral population, and the estimator becomes biased. In contrast, the lineages-remaining estimator is variable at all times but is less affected by bias associated with the period of selection. (For readability, results for the waiting-time estimator are not shown, but they are similar to the lineages-remaining estimator.) The estimates shown are formed from simulated coalescent trees for a sample of 200 chromosomes.

The proportion-of-lineages estimator (Eq. 4) estimates allele frequencies as the proportion of lineages ancestral to the sample carrying each allele. It is expected to perform well under neutrality, and it does here—in the neutral period between the offset of selection and the present, it tracks the true polygenic score closely. During the period of selection, looking backward in time, the proportion-of-lineages estimator strays off target, slowly recovering in the period before the onset of selection. Looking forward in time, it is as if the proportion-of-lineages estimator “anticipates” shifts due to selection. As mentioned in section 3.1.1, the apparent anticipation occurs because if there has been selection between the present and time *t* in the past, then the ancestors of the present-day sample at time *t* are a biased sample from the population at time *t*. For example, if the trait has been selected upward, then the ancestors of a present-day sample will have had high trait values compared with their peers.

The waiting-time estimator (Eq. 11) and the lineages-remaining estimator (Eq. 14) do not rely on an explicit neutrality assumption. Instead, they track the relative passage of coalescent time—measured, roughly, in terms of coalescence events—for each allelic type. These estimators track the rapid change in the polygenic score during the period of selection much more closely than the proportion-of-lineages estimator. (Only the lineages-remaining estimator is shown in Figure 3.) At the same time, these estimators rely on a highly stochastic signal—in the case of the waiting-time estimator (with *ℓ* = 1), single coalescence events—and they are noisier than the proportion-of-lineages estimator as a result.

The patterns seen in Figure 3 figure reflect the performance of the methods over many simulations, as detailed in the next subsections.

### 4.1 Estimator performance: bias and mean squared error

Figure 4 shows bias and mean squared error (MSE) of our estimators of the historical polygenic score across three scenarios—one in which the trait has evolved neutrally, one in which there has been recent directional selection on the trait, and a third in which there has been directional selection on the trait in the past but the trait has evolved neutrally recently. These estimators are also compared with a “straight-line” estimator—a straight line that goes from the present value to the ancestral state (i.e. all derived allele frequencies zero) in two coalescent units. In the neutral case, none of the estimators show marked bias, and the proportion-of-lineages estimator has the lowest variance (and thus lowest MSE). Estimators formed from trees reconstructed by RENT+ (dashed lines) rather than the true trees (solid lines) are noisier, and they do not outperform the straight-line estimator under neutrality.

**Figure 4:**
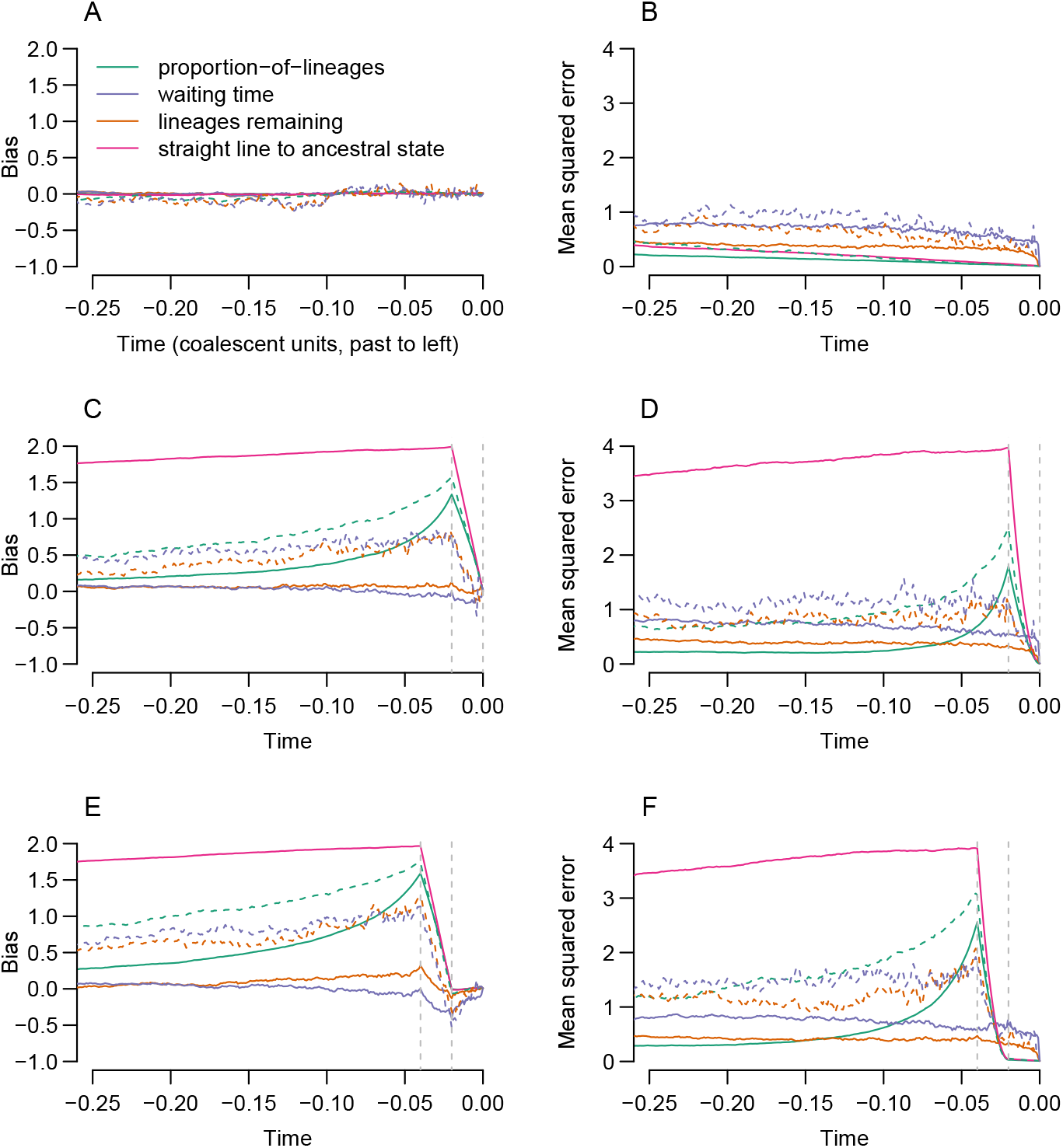
Bias and mean squared error (MSE) of the proposed estimators. For the estimates computed from true coalescent trees (solid lines), 1,000 simulations were performed; RENT+ was used to reconstruct trees for 100 of these trials (dashed lines). In each trial, the polygenic scores were formed from 100 loci. The polygenic scores either evolved neutrally (A-B), or with selection leading to an approximate two-standard deviation shift in mean polygenic score (in present-day units), either occuring from .02 coalescent units ago to the present (C-D), or from .04 to .02 coalescent units ago (E-F).

In the presence of selection, the proportion-of-lineages estimator is badly biased, and the severity of the bias increases during the interval of selection (looking backward in time). The waiting-time and lineages-remaining estimators are less strongly biased in the presence of selection, and they achieve similar MSEs under selection and neutrality. Again, estimators formed from RENT+ trees perform worse than estimators formed from the true trees, but in the presence of selection, they outperform the straight-line estimator.

### 4.2 Interval estimation: coverage probabilities

Figure 5 shows the coverage probabilities of nominal 95% confidence intervals formed on the basis of the proportion-of-lineages, waiting-time, and lineages-remaining estimators and their (approximate) variances.

**Figure 5:**
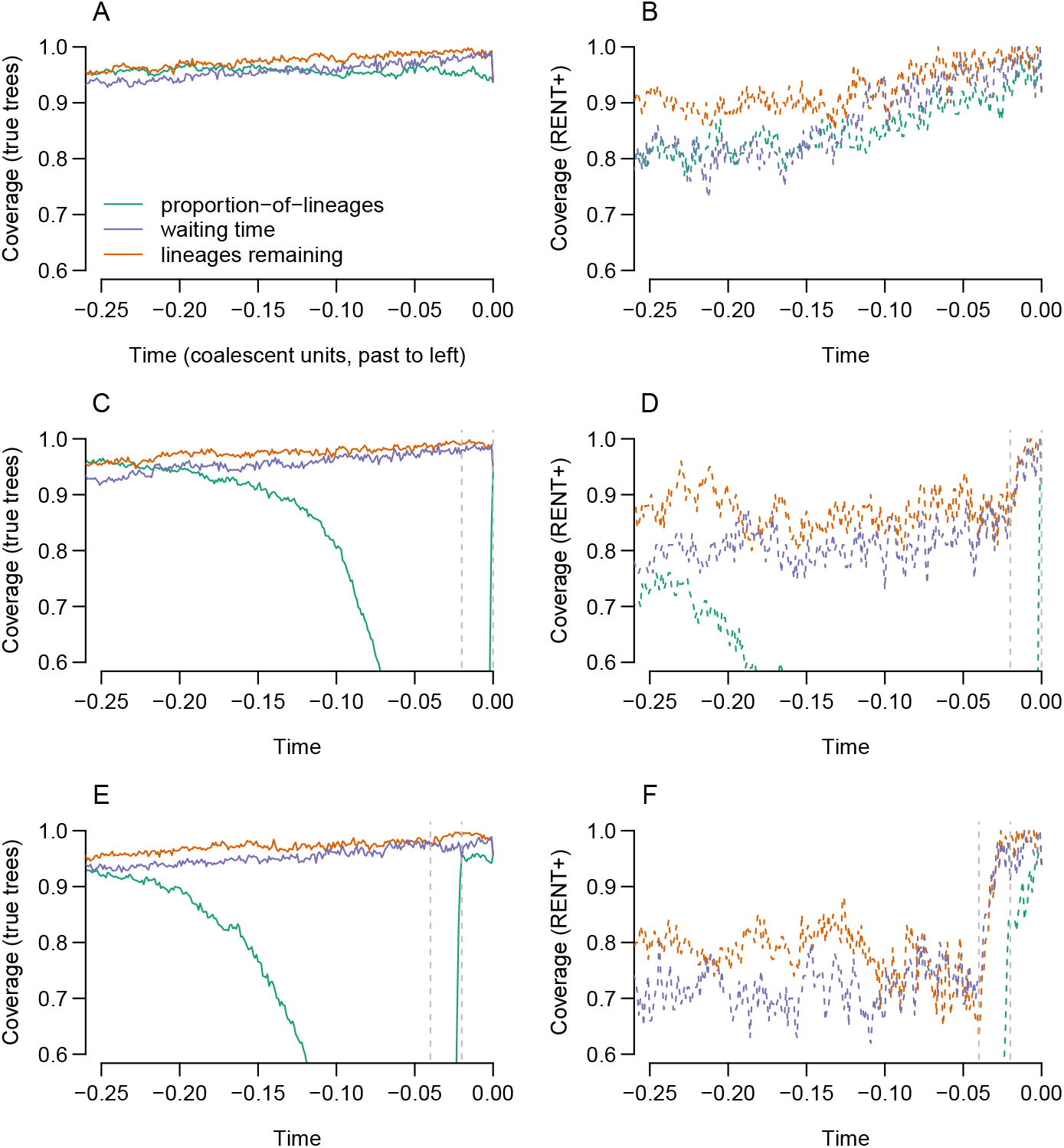
Confidence-interval coverage for nominal 95% confidence intervals based on the proposed estimators. All confidence intervals were formed assuming an approximately normal distribution for the estimator, adding ±1.96 standard errors to the estimate. Standard errors were computed by taking the square root of the approximate variance of each estimator. Coverage probabilities are based on 1,000 simulations (for true trees, figures A, C, E) or 100 simulations (for RENT+ trees, figures B, D, F). Simulations were conducted under either neutrality (A-B), an approximate two-standard-deviation shift over the last .02 coalescent units (C-D), or a two-standard-deviation shift from .04 to .02 coalescent units ago (E-F).

Under neutrality, and when using the true trees, all confidence intervals have approximately the correct coverage, though coverage for confidence intervals for both the lineages-remaining and waiting-time estimators decays further into the past. The decay of the coverage probability far in the past makes sense for the lineages-remaining and waiting-time estimators—both these estimators implicitly assume that the number of carriers of each allele in the population remains constant between coalescent events. This assumption may be a reasonable approximation in the recent past, when coalescent events are frequent, but become untenable in the distant past, when coalescence times are longer.

Confidence intervals computed on the basis of RENT+ trees only achieve the nominal coverage in the very recent past and become anticonservative further back in time. This behavior is expected; the variances we use incorporate stochasticity in the coalescent process but do not account for randomness arising from errors in tree estimation. (Stochasticity in the coalescent process has been called “coalescent error” and contrasted with randomness from errors in tree estimation, or “phylogenetic error” (Ho and Shapiro, 2011).)

Under selection, confidence intervals from the proportion-of-lineages estimator have very low coverage—this arises from the bias documented in Figure 4. The coverage probabilities of the waiting-time and lineages-remaining estimators are less changed by selection.

### 4.3 Power of *T_X_*

We assessed the performance of the *T_X_* statistic (Eq. 16) as a test statistic for detecting selection in the simulations shown in Figures 4–5. We constructed the test statistic from the allele-frequency estimates produced by each of the proposed estimators and compared it against both the theoretical *χ*^2^ distribution and against a permutation distribution. Table 1 shows the results. Under neutrality (first two rows), comparing *T_X_* against a distribution formed by randomly permuting the effect sizes produces acceptable type I error rates. (There are 100 simulations using RENT+, and the values in Table 1 do not differ significantly from .05.) When the theoretical *χ*^2^ distribution is used, RENT+ type I error rates are unacceptably high, but the type I error rates produced from the true trees are acceptable.

**Table 1:** Power/Type I error (with *α* = .05) of various implementations of the *T_X_* statistic in the simulations shown in Figures 4–5, with 200 chromosomes sampled in the present. Power is not shown for methods with type I error rates ≥ .1. *T_X_* was computed using allele-frequency estimates 0, .01, .02, …, .1 coalescent units before the present.

|  |  | proportion-of-lins |  | waiting-time |  | lins-remaining |  |
| --- | --- | --- | --- | --- | --- | --- | --- |
| | | $\chi^2$ | perm. | $\chi^2$ | perm. | $\chi^2$ | perm. |
| neutral | true trees | .053 | .054 | .030 | .060 | .061 | .064 |
|  | RENT+ | .17 | .08 | .31 | .04 | .11 | .09 |
| sel. .02-now | true trees | 1 | 1 | .033 | .060 | .194 | .222 |
|  | RENT+ | - | .87 | - | .05 | - | .15 |
| sel. .04-.02 | true trees | .969 | .954 | .035 | .065 | .058 | .084 |
|  | RENT+ | - | .37 | - | .08 | - | .02 |

Of the methods with acceptable type I error, tests using the allele frequencies estimated by the proportion-of-lineages estimator have by far the highest power. It may seem paradoxical that the proportion-of-lineages estimator is the best of our estimators at detecting selection, given that the estimated time courses produced by the proportion-of-lineages estimator are biased in the presence of selection. However, in our simulations, the proportion-of-lineages estimator generally moves in the correct direction in the presence of selection, albeit more slowly than it should. In contrast, the other two allele-frequency estimators are highly variable, leading to wide null distributions and decreased power. The proportion-of-lineages estimator can also be thought of as the mean polygenic score among lineages ancestral to the sample, and the test for selection responds for changes in the mean polygenic score of the ancestors that are faster than would be expected under the null hypothesis of neutral evolution.

With the proportion-of-lineages estimator, using true trees unsurprisingly gives better power than RENT+ trees, but RENT+ trees still have substantial power.

In Table 1, power is higher when selection occurs closer to the present. To explore the relationship between the timing of selection and present-day sample size, we conducted additional simulations. In these simulations, we assessed the power of the *T_X_* test (using the proportion-of-lineages allele-frequency estimates from true trees, and comparing with a permutation distribution) to detect an approximate one-standard-deviation shift in the population-mean polygenic score. We varied the timing of the shift and the present-day sample size. Figure 6 shows the results. For detecting selection close to the present, power increases with sample size. However, for selection further in the past, power reduces to the type I error rate, regardless of the present-day sample size. This is because power to detect selection depends on unusual coalescent times during the period of selection, and by .1 coalescent units in the past, most coalescent events have already occurred, even in large samples. For example, even extremely large present-day samples have, in expectation, ~ 200.5 ancestors tracing back 0.01 coalescent units in the past and ~ 20.5 ancestors 0.1 coalescent units in the past (Maruvka et al., 2011; Jewett and Rosenberg, 2014). Thus, it will likely be impossible to detect all but the strongest selective events by their signatures in coalescent trees if they are over 0.1 coalescent units in the past. *T_X_*’s power to detect selection up to ~ .02 − .04 coalescent units into the past represents an extension of the SDS statistic (Field et al., 2016, Figure S6), which has excellent power in the very recent past but very little power beyond the expected length of a terminal branch (2/*n* in coalescent units, where *n* is the present-day sample size (Fu and Li, 1993)). In appendix F, we show empirical power for a test statistic analogous to SDS computed from the lengths of the terminal branches. This SDS analogue has similar power to *T_X_* near the present, but its power decays more rapidly for selection further in the past.

**Figure 6:**
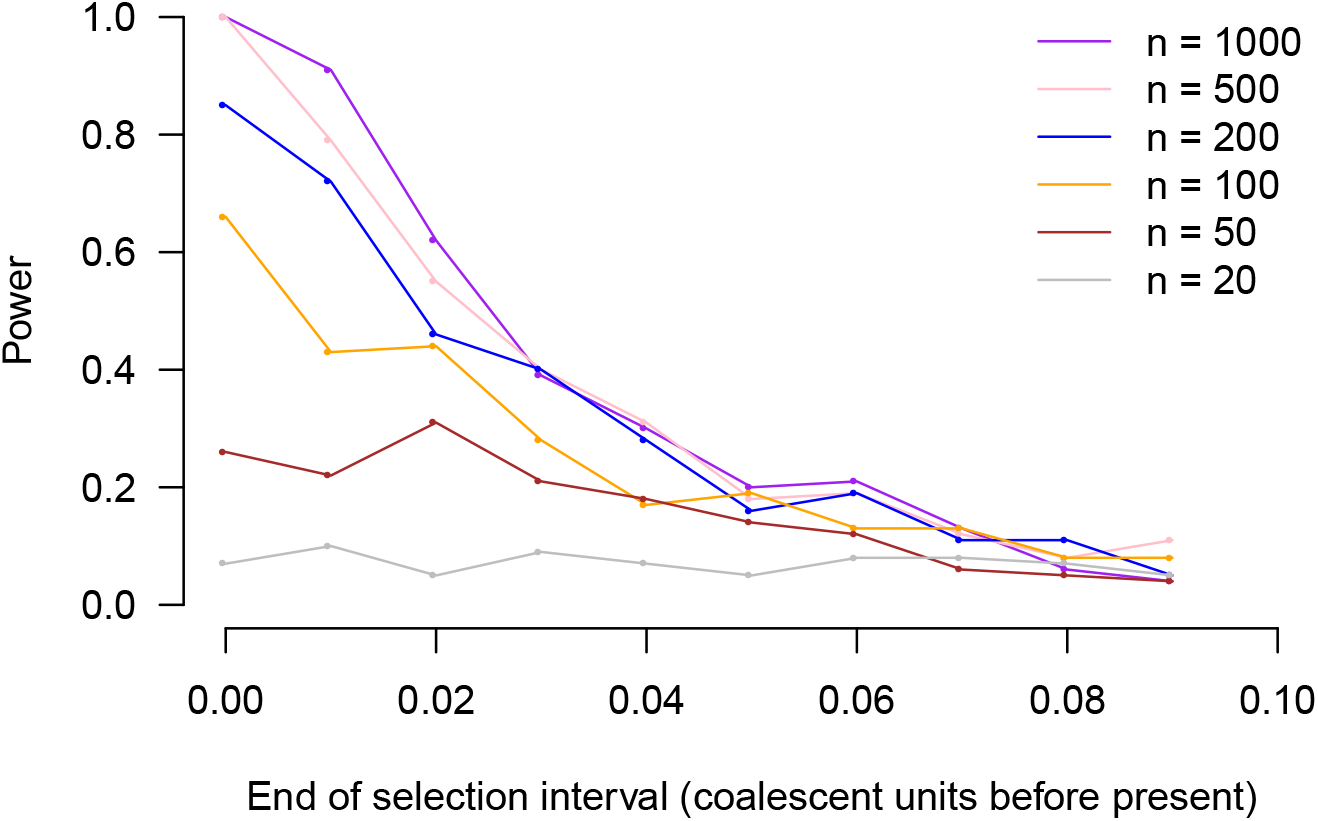
Power of the *T_X_* test as a function of timing of selection and present-day sample size. Power is shown on the vertical axis. The horizontal axis shows the timing of a pulse of selection lasting .005 units and resulting in an approximate one-standard-deviation shift in the population-average polygenic score (in present-day standard deviations). Each point is based on power from 100 simulations, using the proportion-of-lineages estimator on the true trees, and using a null distribution obtained by randomly permuting locus effect sizes.

## 5 Empirical application: Human height

We applied our proposed estimators to human polygenic scores for height. Genetic variation within Europe related to human height has been studied by many investigators interested in polygenic selection (Turchin et al., 2012; Berg and Coop, 2014; Robinson et al., 2015; Field et al., 2016; Berg et al., 2017; Racimo et al., 2018; Uricchio et al., 2018). A recent pair of papers compared the results produced by existing tests for polygenic selection when applied to human height using genome-wide association study (GWAS) effect sizes from two different studies (Berg et al., 2018; Sohail et al., 2018). Most previous work has used GWAS effect sizes from the GIANT consortium (Wood et al., 2014), whereas the new work uses GWAS effect sizes from the larger and presumably less structured UK Biobank sample (Sudlow et al., 2015). Tests for polygenic selection on height provide much less evidence for selection when UK Biobank effect sizes are used than when effect sizes from GIANT are used. One possible explanation is that GIANT effect sizes are contaminated by some degree of population stratification.

In Figure 7, we show estimated population-mean polygenic score time courses among populations ancestral to the GBR (British in England and Scotland) subsample of the 1000 Genomes Project (1000 Genomes Project Consortium, 2012). In the top panel, GIANT effect sizes are used, and in the bottom panel, UK Biobank effect sizes are used. Polygenic scores were constructed by taking the top locus in each of ~ 1700 approximately independent genetic regions. (These polygenic scores are identical to those used in Berg et al. (2018); our “UK Biobank” is their “UKB-GB.”) Coalescent trees for these loci were estimated in RENT+. (Details in Appendix G.)

**Figure 7:**
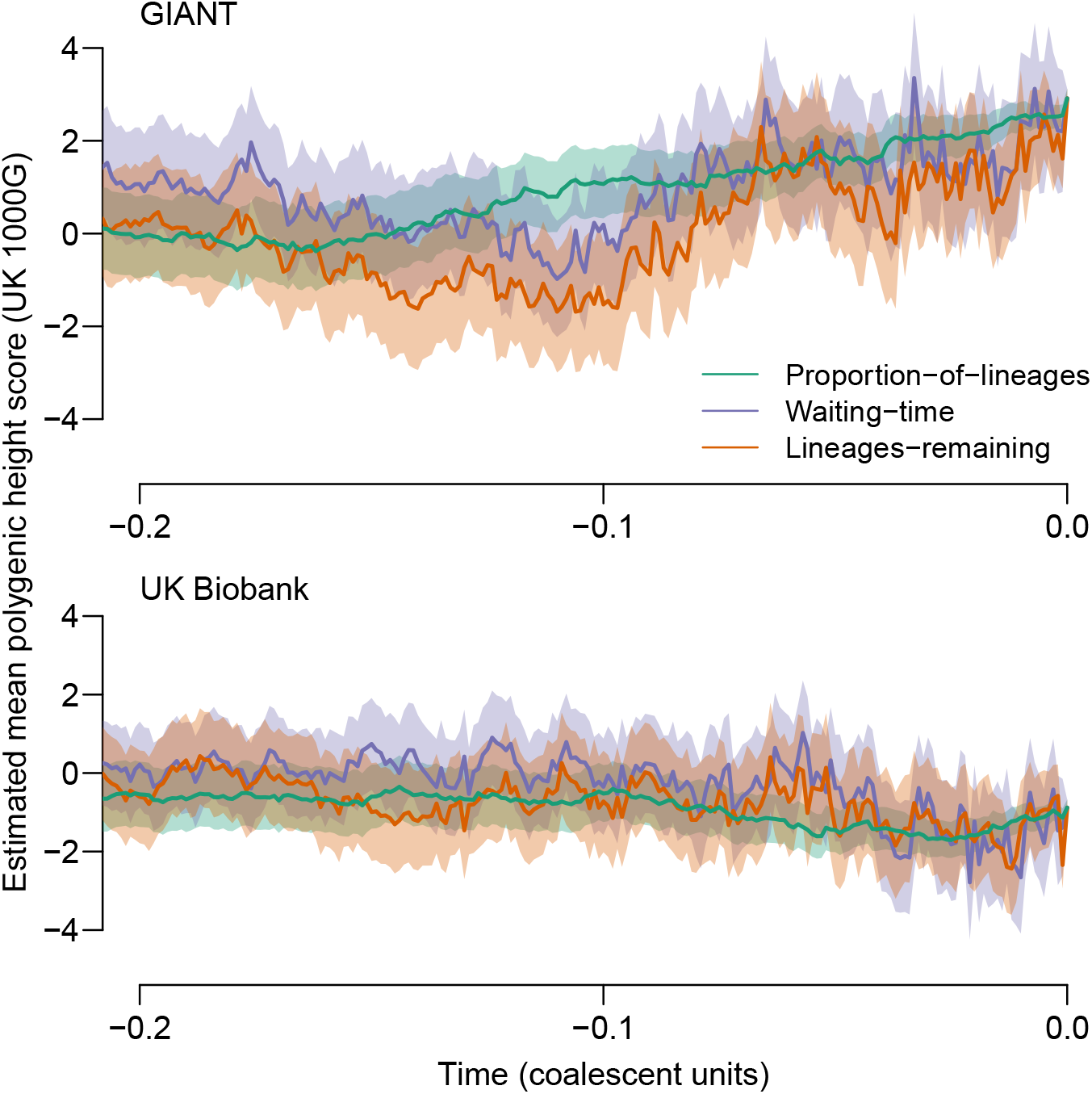
Estimated population-mean polygenic scores for height in the population ancestral to the GBR subset of the 1000 Genomes project (91 genomes). Time is displayed in approximate coalescent units, defined such that the mean across loci of the time (measured in mutations) to the most recent common ancestor is 2. To construct the polygenic scores, the top locus in each of ~ 1700 approximately unlinked genomic regions was chosen. (See Appendix G for more information.) The top panel shows the results when effect sizes estimated by GIANT are used; the bottom panel shows analogous results when UK Biobank effect-size estimates are used. In both panels, the effect sizes are scaled so that the standard deviation of the polygenic score is 1 in the present, assuming linkage equilibrium among SNPs.

When the time courses are constructed using GIANT, all three estimators suggest that the population-mean polygenic score for height has increased in the recent past. Using the proportion-of-lineages approach, an increase of ~ 3 present-day polygenic score standard deviations is estimated. In contrast, time courses estimated using the UK Biobank effect sizes show little apparent change in the recent past—the proportion-of-lineages estimator suggests a recent decrease of ~ .34 standard deviations.

Further, when the change from the present to the most recent timepoint (.001 units in RENT+) is assessed by the *T_X_* test, both sets of effect sizes yield some evidence of selection, but the evidence is stronger with GIANT effect sizes than with UK Biobank effect sizes. Specifically, with GIANT, *T_X_*(1) = 14.9, *p* = .0005 from 10,000 permutations, and with UK Biobank, *T_X_* (1) = 7.0, *p* = .0112. (We do not claim that the difference between the *T_X_* values across the datasets is itself significant (Gelman and Stern, 2006), merely that the pattern of weakened evidence in the UK Biobank matches that observed by recent work (Berg et al., 2018; Sohail et al., 2018).) However, using both datasets, the evidence for selection is limited to periods very close to the present. If the same set of times evaluated in the simulations is evaluated for the height polygenic scores (approximate coalescent times 0, .01, .02, …, .1 before the present), neither polygenic score time course provides evidence for selection (*p* ≈ .2 in both cases). GIANT effect sizes produced much lower p values for recent selection on height in the UK in a recent paper (Field et al., 2016), but that work used a sample of 3195 genomes, whereas the GBR subset of the 1000 Genomes sample contains only 91 genomes.

Thus, our estimators broadly recapitulate the pattern of other methods for detecting polygenic selection, finding evidence suggestive of selection when GIANT effect sizes are used but much weaker evidence when UK Biobank effect sizes are used.

## 6 Discussion

We have proposed a set of estimators and tests for population-mean polygenic scores over time, given (additive) effect sizes for a trait at independent trait-associated loci, and coalescent trees for the trait-associated loci. Estimation of the population-mean polygenic score time course is most effective when the trait (and its associated loci) evolve neutrally, and the ancestors of the sample are representative of the ancestral population. When the trait has been under selection, estimation is still possible, but the estimates obtained are noisier. Tests for polygenic selection that are based on coalescent trees have the potential to be powerful in the recent past.

In terms of practical applications, we have produced one estimator that produces good estimates of population-mean polygenic score time courses under neutrality and that is also well-powered to detect departures from neutrality (the proportion-of-lineages estimator). The other two estimators are less biased by selection, but they are variable and less useful for detecting selection. At this writing, one sensible procedure for fitting these methods to data would be to form initial estimates using the proportion-of-lineages estimator and test them for selection using the *T_X_* statistic. If the test suggests selection, then the ancestors of the sample may not be representative of the ancient population, and polygenic-score time courses from the proportion-of-lineages estimator may be biased. In that case, the waiting-time or lineages-remaining estimators might be applied.

These methods add to a set of methods that use GWAS information to study the history of complex traits (Berg and Coop, 2014; Field et al., 2016; Berg et al., 2017; Racimo et al., 2018; Uricchio et al., 2018). Many of these methods have been applied to human height, and our methods produce similar conclusions when applied to the same data (Berg et al., 2018; Racimo et al., 2018; Sohail et al., 2018; Uricchio et al., 2018). Our methods add to previous work by estimating the historical time courses of mean polygenic scores and by leveraging ancestral recombination graphs.

As with other population-genetic methods for studying polygenic traits, results from our methods are accompanied by many qualifiers to interpretation (Novembre and Barton, 2018).

In general, estimates arising from these methods should not be viewed as necessarily reflecting the historical time course of trait values within a population. Rather, they reflect the history of a function of the genotype that encodes present-day associations between genotype and phenotype. Real-world estimates of effect size will be subject to noise in estimation and possibly bias due to stratification. (Even small amounts of stratification can seriously mislead tests for selection (Berg et al., 2018; Sohail et al., 2018)) Particularly in case-control studies, ascertainment biases may also lead to confounding between the evolutionary status of an allele (i.e. derived or ancestral status) and power to detect trait associations (Chan et al., 2014). Further, the genetic architecture may change across the time period over which estimates are made, for example because of changes in linkage disequilibrium between tag loci and causal loci (Martin et al., 2017), or because loci that explained trait variation in the past have since fixed or been lost. And, importantly, changes in the environment may drive changes in mean levels of the trait that either amplify or oppose changes in population-mean polygenic scores, either via their direct effects or via gene-environment interactions.

Beyond these general caveats, the methods we propose here have several limitations that suggest directions for future work. The first three limitations concern outstanding statistical issues. First, the polygenic scores we estimate here are linear combinations of effect sizes estimated under additive models. Our variance estimates also assume that the loci incorporated in the polygenic score are in linkage equilibrium. Because our estimators work by estimating historical allele frequencies at the loci contributing to the polygenic score, they can in principle be adapted to estimate any function of allele frequencies, including trait predictions that account for dominance, epistasis, and linkage among loci. However, the strategies we use here for variance estimation and hypothesis testing may need to be modified for more general functions of allele frequencies. Second, any applications of these methods to real data will entail noise that is not accounted for by the variance estimates we propose. In particular, effect sizes will be estimated with error, and so will the coalescent trees for sites included in the polygenic score. It will be important to incorporate these sources of variance in future estimates of sampling variation. Third, as suggested in section 3, the waiting-time and lineages-remaining estimators implicitly contain smoothing parameters. Fully characterizing the effects of these smoothing parameters and of alternative smoothing strategies—such as those used to smooth coalescent-based estimates of population-size history (Drummond et al., 2005; Minin et al., 2008)—will reveal the potential of these estimators, which have high variance in the forms in which they are used here.

The next three possible extensions are suggested by biological applications and by the coalescent framework in which we work. First, the theory we develop here is for a single population, but our setting within a coalescent framework suggests the possibility for extension to multiple populations, perhaps by developing multivariate analogues of our statistics within a coalescent-with-migration framework (Kaplan et al., 1991). Similarly, whereas we work with polygenic scores for a single trait, our methods can be extended to consider polygenic scores for multiple, correlated traits. In a similar vein, Berg et. al (2017) have recently extended the *Q_X_* statistic to multiple correlated traits, drawing inspiration from the framework of Lande and Arnold (1983). Working with multiple traits will allow us to distinguish hypotheses in which a trait is directly subject to selection from hypotheses in which a correlated trait is the target of selection. Finally, because the coalescent framework explicitly represents the evolution of the sample backward in time, it will be productive to incorporate ancient samples.

The methods we propose are promising in part because they capitalize on illuminating descriptions of genetic variation that will become increasingly widely available. Ancestral recombination graphs encode all the coalescence and recombination events reflected in the present-day sample, and thus are richly informative about the history of the sample’s ancestors. Statistics computed from these ARGs have the potential to capture all the information about an allele’s frequency time course that is available in a present-day sample. Approaches to traits that are based on sample ARGs will improve with the development of our understanding of the architecture of complex traits and of our ability to reconstruct ARGs.

## 7 Acknowledgments

We thank members of the Coop lab and Arbel Harpak for useful discussions, Jeremy Berg for providing the height loci and effect sizes used in Berg et al. (2018), and Sajad Mirzaei for support with RENT+ software. The authors acknowledge support from NIH (R01-GM108779) and NSF (1262327 and 1353380).

### A The relationship between coalescent rates and phenotypic selection

In this appendix, we describe the consequences of directional and stabilizing selection on a trait for coalescence rates at loci associated with the trait.

We begin by considering the general effects of selection on coalescence before considering the specific cases of directional and stabilizing selection. If the frequency of the allele of interest at locus *i* at time *t* is *p_i_*(*t*) in a population of 2*N* chromosomes, then the probability of any one of 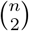 pairs coalescing in the preceding generation is

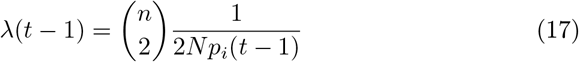

Hudson and Kaplan (1988). Writing the frequency in the preceding generation as *p_i_*(*t* − 1) = *p_i_*(*t*) − Δ_*i*_(*t*), then

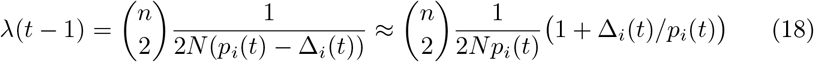

assuming that Δ_*i*_(*t*) is small so that ignore higher-order terms in the Taylor expansion can be ignored. We can decompose the change in frequency from the preceding generation into 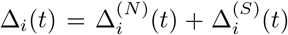, the contributions of random genetic drift (*N*) and selection (*S*) respectively. Then taking the expectation over the frequency in the preceding generation,

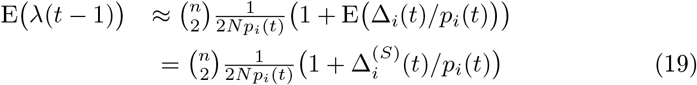

because 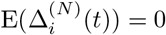 and 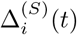 is the deterministic change due to selection.

If the allele of interest affects a trait that is under selection, then the form of 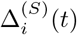 depends on the allele’s role in the trait’s architecture and on the specifics of selection on the trait. The simplest case is one in which directional selection acts on a phenotype whose genetic architecture is purely additive. Denote the selection gradient—that is, the slope of fitness regressed on phenotype—as *α*. If the effect size of the *i^th^* locus on the phenotype is *β_i_*, and the loci that affect the phenotype are unlinked, then

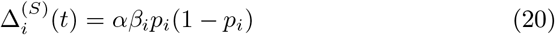

(Charlesworth and Charlesworth, 2010, Eq. 3.17). Therefore, the expected coalescent rate over the possible allele frequency changes in the preceding generation is

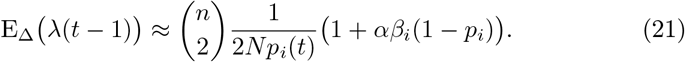

Therefore, recent directional selection increases the coalescence rate among alleles that move the phenotype in the direction in which selection acts. For example, if larger trait values have been selected recently, then the coalescence rate will be higher among alleles associated with larger values of the trait and lower among alleles associated with smaller values of the trait. This is true regardless of the frequency of the allele.

If the phenotype is subject to both directional selection and stabilizing selection, then the allele frequency matters. For example, one common and reasonably general model incorporating directional and stabilizing selection is the quadratic optimum model, where the fitness of an individual with phenotype *G* is *W*(*G*) = 1 − *s*(*P_O_* − *G*)^2^. There is directional selection on the phenotype when the population mean (Ḡ) deviates from the optimum *δ* = *P_O_ − Ḡ* ≠ 0. (When *δ* > 0, the population mean is below the optimum, and selection will favor larger trait values.) Under this model,

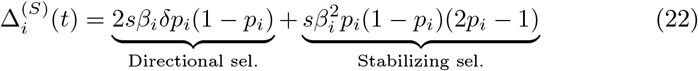

see Barton (1986); Bürger (2000) and Simons et al. (2018) for a recent discussion. Note the similarity of the directional selection term—with the selection gradient now controlled by *sδ*— to Eq. (20), resulting in a similar coalescent rates to Eq. (21). The Δ term for stabilizing selection matches that of a disruptive-selection model with an unstable equilibrium at 1/2 − *δ*/*β_i_*.

Incorporating both directional and stabilizing selection, the expected coales-cent rate is approximately

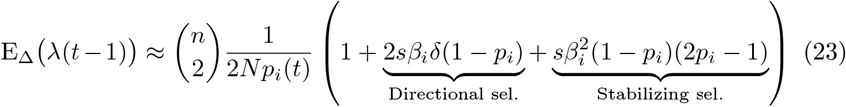

The stabilizing selection term decreases the coalescent rate of whichever allele has frequency less than 1/2. The average coalescent rate among “up” alleles (*β_i_* > 0) depends on the value of E_Δ_(λ(*t* − 1)) averaged over the frequency distribution of *p_i_*. Thus, when directional selection acts on the phenotype, we again expect an increased coalescence rate among alleles whose effects have been favored. If the population is at the optimum, i.e. *δ* = 0, then averaging across loci there is no net effect on the coalescent rates if there is a symmetric distribution of *p_i_* around 1/2 (across all effect sizes). This symmetric distribution will result when the population is at equilibrium and there is no mutational bias (i.e. there is no difference in the mutational input of up and down alleles).

### B A Bayesian view of the proportion-of-lineages estimator

A Bayesian view of the estimator proposed in Eq. 4 relies on connections between neutral diffusion processes and coalescent processes, reviewed by Tavaré (1984).

Under a neutral diffusion model without mutation, the probability that an allele frequency changes from *p_i_*(0) = *p* to *p_i_*(*t*) = *x* in time *t* can be written as (Tavaré 1984, Eq. 7.18)

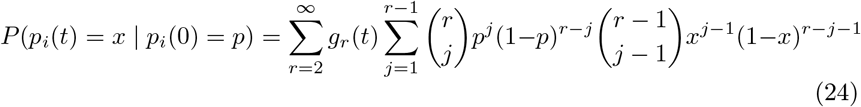

where *g_r_*(*t*) is the probability that *r* lineages of an initially infinite number of lineages survive to time *t*. One way to interpret Eq. 24 is that if *r* lineages survive to time *t*, then the number *j* of lineages carrying an allele at time *t* is a Binomial(*r,p*) random variable. Then, conditional on *j*, the frequency *p_i_*(*t*) has a Beta(*j, r − j*) distribution. One way to interpret the Beta(*j, r − j*) distribution of *p_i_*(*t*) given *j, r*, and *p* is as a posterior distribution resulting from multiplying an improper prior proportional to *x*^-1^(1 − *x*)^-1^—which is proportional to the limit as the mutation rates go to 0 of the stationary distribution of the neutral diffusion with mutation (Ewens, 2004, Eq. 5.70)—by the probability mass function of a Binomial(*r,p*) distribution evaluated at *j*.

Thus, conditional on the number of lineages at time 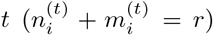, the number of lineages carrying the allele of interest 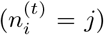, and the allele frequency in the present (*p*), we have

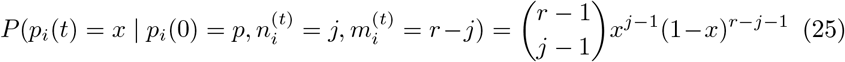

Eq. 25 suggests the posterior mean of *p_i_*(*t*) as an estimate of the frequency (when *r* ≥ 2 and *j* ≥ 1), which is equal to Eq. 4. The posterior variance is

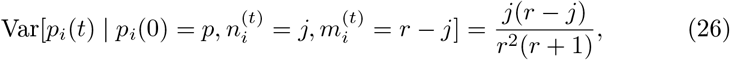

which differs from the binomial sampling variance in Eq. 5 by a factor of *r*/(*r* + 1).

### C Mathematical details for the waiting-time and lineages-remaining estimators

#### C.1 Taylor-series expansions for expectations and variances of ratios

Both the waiting-time and lineages-remaining estimators take the form

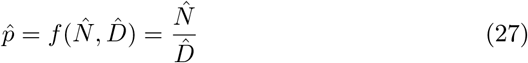

where 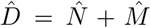, and 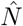 and 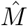 are population-size estimators that are independent of each other. We assume that 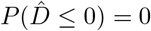. Eq. 27 is equivalent to Eq. 7 from the main text, with the subscripts and time notation removed for compactness.

In this subsection we present general Taylor-series approximations for the expectation and variance of the estimator in Eq. 27 in terms of the expectation and variance of 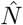 and 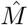. Similar presentations are available in other references (Stuart and Ord, 1987, sec. 10.5-10.6), but we present the argument here for completeness.

The first-order Taylor expansion of Eq. 27 evaluated at 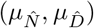 is

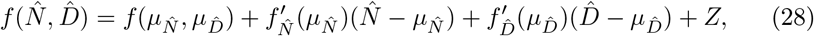

where 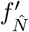 and 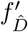 are the partial derivatives of *f*, and *Z* represents the contribution of higher-order terms. The expectation of 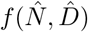 is therefore

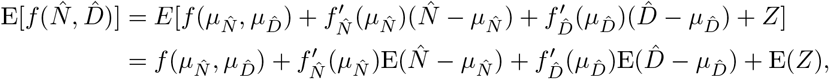

with the second step coming from the linearity of expectation and the fact that 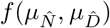 is not random. Choosing 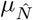 and 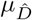 to be equal to the expectations of 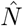 and 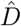 gives

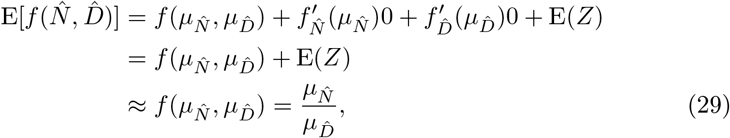

where the approximation depends on the higher-order terms represented by *Z* being small. Equation 29 justifies the claim in the main text that the waiting-time estimator is approximately unbiased (under the unrealistic assumption that the number of carriers of each allele is constant between coalescent events).

To obtain the approximate variance of the estimator in Eq. 27, we use Eq. 29 and note that

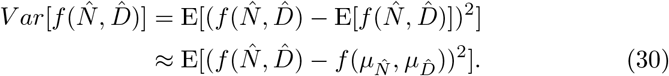

Substituting the first-order Taylor expansion in Eq. 28 for 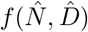, ignoring the higher-order terms represented by *Z*, gives

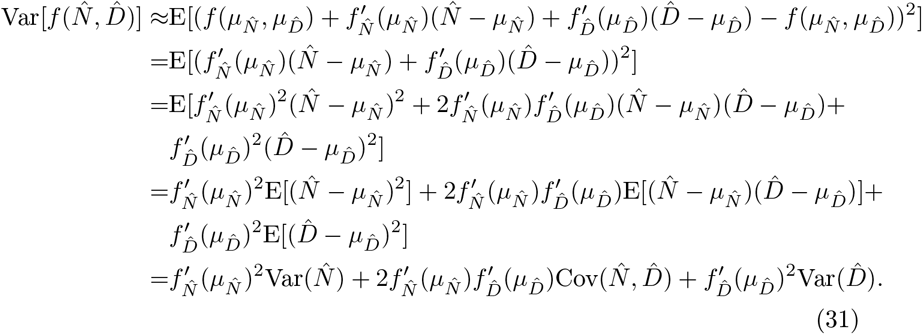

The last step comes from the definitions of variance and covariance and the fact that we choose 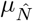 and 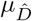 to be equal to the expectations of 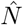 and 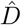. In our case, 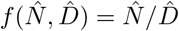, implying that 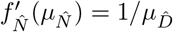 and 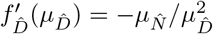. Plugging these values into Eq. 31 gives

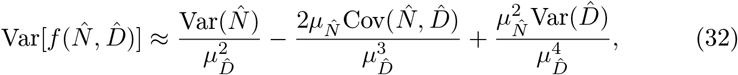

with the useful alternative form

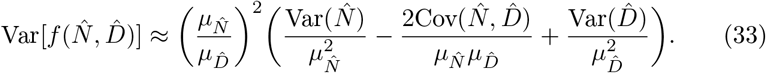

Finally, in our setting, 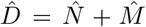, and 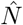 and 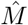 are independent, giving 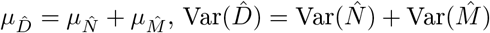 and 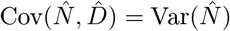. Using these identities in Eq. 33 gives

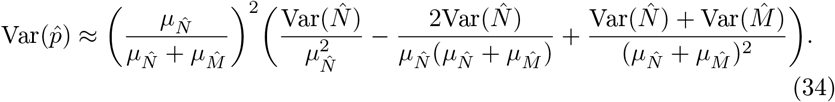

#### C.2 Approximate variance of the waiting-time estimator

As noted in the main text, the waiting-time estimator has the form

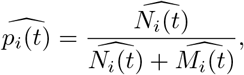

where

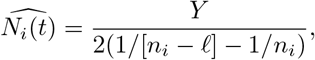

*Y* is the waiting time associated with a set of *ℓ* coalescent events on the allele of interest’s background that envelop time *t*, and *n_i_* is the number of allele-of-interest lineages that exist before any of the *ℓ* coalescent events have occurred. 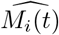 is analogous, substituting in a waiting time and number of lineages from the other allele’s background, and possibly a different value of *ℓ*.

It is also noted in the main text that the waiting time *Y* has variance

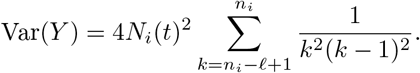

Thus, the variance of 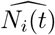 is

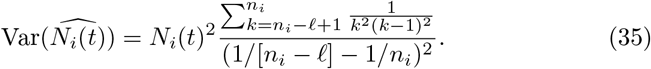

If *ℓ* =1, Eq. 35 reduces to 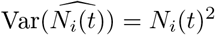. If *ℓ* << *n_i_*, then

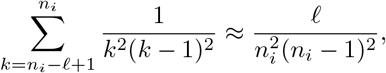

and 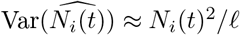.

By Eq. 34 and the fact that 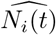 and 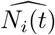 are unbiased, the approximate variance of the waiting-time estimator is

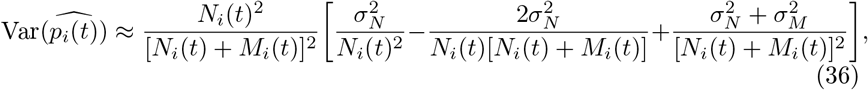

where 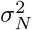 and 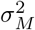 are the variances of *N_i_*(*t*) and *M_i_*(*t*), respectively. If *ℓ* =1 is used to estimated both *N_i_*(*t*) and *M_i_*(*t*), then 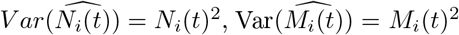, and Eq. 36 reduces to

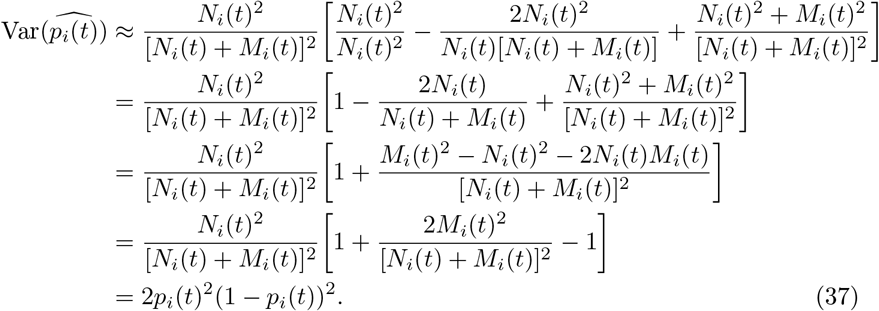

If *N_i_*(*t*) and *M_i_*(*t*) are estimated using the same value of *ℓ*, and *ℓ* is much smaller than the starting number of lineages for both loci, so that 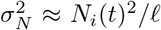 and 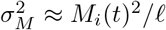, then Eq. 36 can be rewritten as

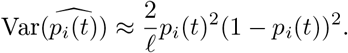

#### C.3 Approximate variance of the lineages-remaining estimator

In the main text, we propose to estimate *N_i_*(*t*) as

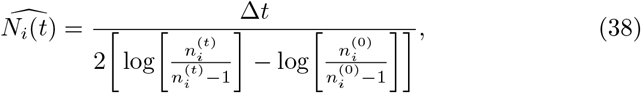

where Δ*t* is the number of generations elapsed between two ends of a predefined time interval, *n_i_*(0) is the number of lineages of the allele of interest at the end of the time interval closer to the present, and *n_i_*(*t*) is the number of lineages of the allele of interest at the end of the interval further into the past. The only random component of 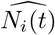 is *n_i_*(*t*), which depends on the number of coalescent events that occur during the interval. This estimator of *N_i_*(*t*) is leads to an estimator for *p_i_*(*t*), the allele frequency of interest,

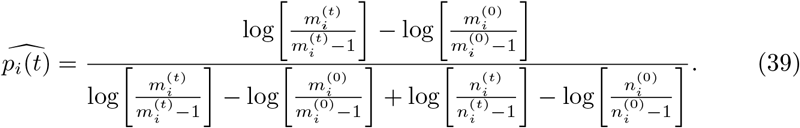

(The expression in Eq. 39 is the same as in Eq. 14.)

To obtain an approximate sampling variance for 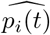, we use Eq. 34, making the substitutions

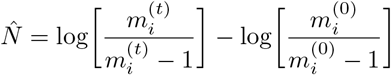

and

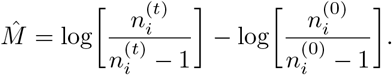

To use Eq. 34, we need (at least approximate) expectations and variances for both these terms. To compute an approximate expectation, we replace 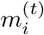 and 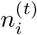 with their expectations. Taking the case of 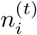, notice that by Eq. 12,

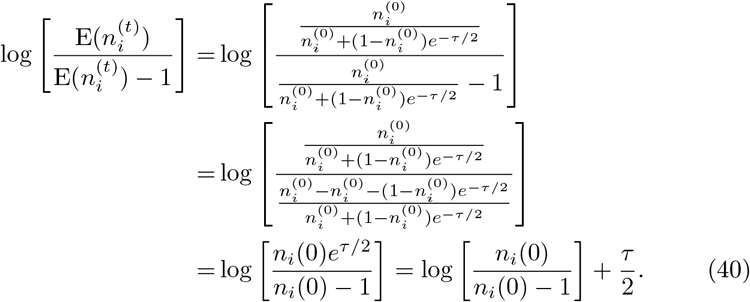

In Eq. 40, *τ* represents the coalescent time passed on the allele of interest’s background, or Δ*t*/*N_i_*(*t*), and so

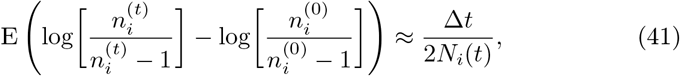

and analogously for 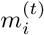,

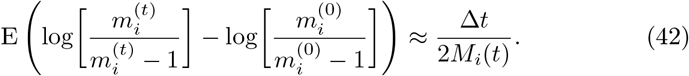

Next we compute the approximate variances of these terms. Because 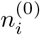 is a constant,

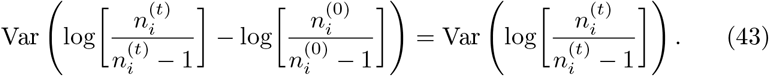

By a first-order Taylor approximation, the variance is approximately

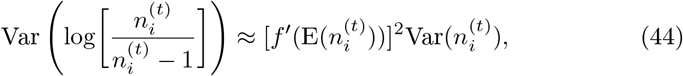

where 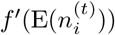 is the derivative of the right side of Eq. 43 with respect to 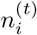, evaluated at 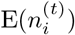. (The argument for this approximation is entirely parallel to the one in Eq. 31, but for a function of a single random variable.) The derivative of the right side of Eq. 43 with respect to 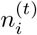, evaluated at 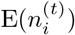, is

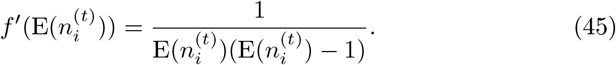

To write the approximate variance of 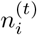, we use the asymptotic variance derived by Griffiths (1984, Eq. 5), using the version for *β* = 0 and *α* < ∞ (see also Chen and Chen (2013)). To change the expression into our notation, we make the replacements 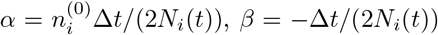 (because we assume only a single mutation distinguishing the two allelic types, so *θ* = 0 conditional on that mutation), and 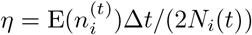 (by Griffiths’ Eq. 4). Doing so and simplifying yields

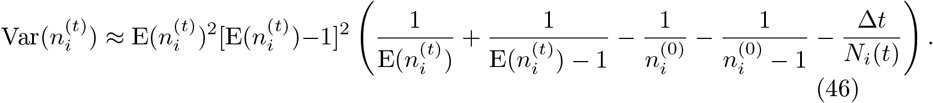

Plugging the expressions for 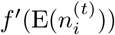 and 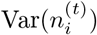 into Eq. 44 gives an approximate variance,

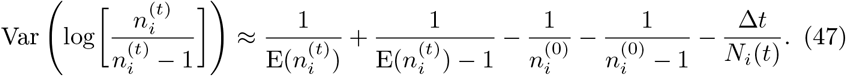

Analogously, for the 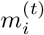 term,

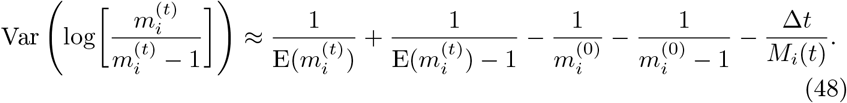

Now, to obtain an approximate variance for 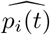, we plug the expressions in Eqs. 41, 42, 47, and 48 into Eq. 34. Doing so gives the expression

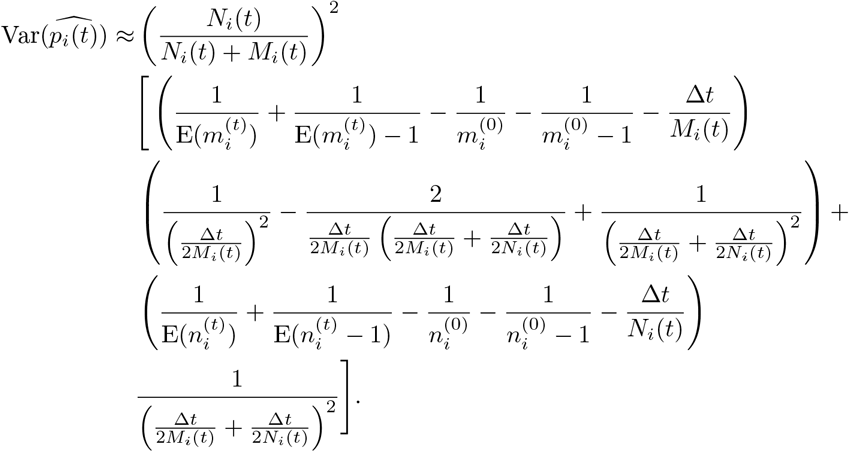

Noticing that *p_i_*(*t*) = *N_i_*(*t*)/[*N_i_*(*t*) + *M_i_*(*t*)], expanding the products in the denominators, and factoring out common terms gives

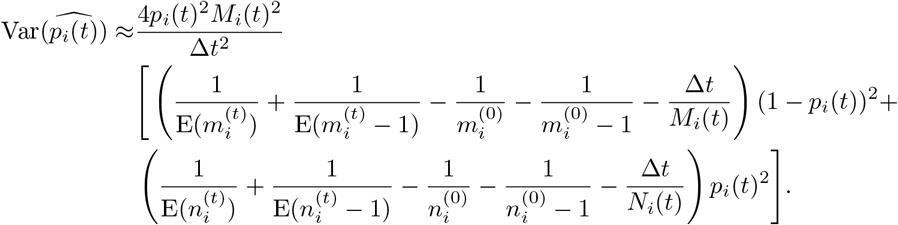

Finally, noticing that *N_i_*(*t*) = *p_i_*(*t*)(*N_i_*(*t*)+*M_i_*(*t*)) and *M_i_*(*t*) = (1 − *p_i_*(*t*))(*N_i_*(*t*) + *M_i_*(*t*)) and defining the coalescent time passed with respect to the whole sample, *τ_s_* = Δ*t*(*N_i_*(*t*) + *M_i_*(*t*)), gives

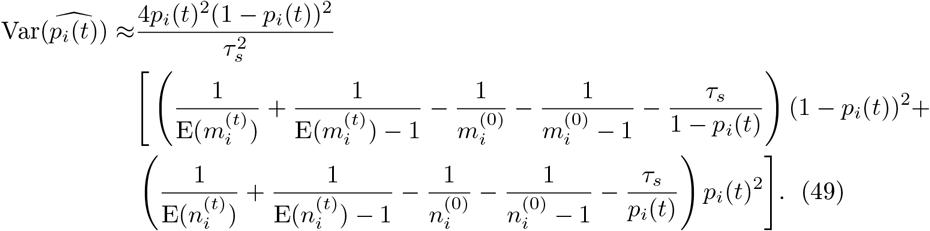

To estimate the approximate variance in Eq. 49, we replace *p_i_*(*t*) with its estimated value from the lineages-remaining estimator, *τ_s_* with the (known or estimated) coalescent time elapsed between timepoints, and the expectations of 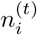 and 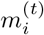 with their realized values.

The approximate variance in Eq. 49 relies on an asymptotic variance of the number of ancestral lineages from Griffiths (Griffiths, 1984). In practice, Eq. 49 gives variance estimates that are much too large when the number of lineages are small and the time passed is short. For example, with 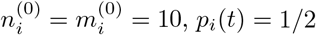, and *τ_s_* = .001, Eq. 49 produces ~ 1.4. Because 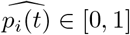, its maximum variance is .25. Our strategy is to use the minimum of Eq. 49 and the estimated variance of the waiting-time estimator with 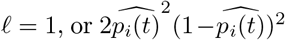 (Eq. 37). The rationale is that 2*p_i_*(*t*)^2^(1 − *p_i_*(*t*))^2^ represents the variance obtained when one estimates *p_i_*(*t*) from only one coalescent event on each background—the lineages-remaining estimator is always based on at least one coalescent event on each background.

#### C.4 Computing the lineages-remaining estimator for edge cases

If no coalescences have occurred in the interval and 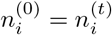, then the estimator in Eq. 13 is undefined. In this case, we compute the estimator for a larger value of Δ*t* in which at least one coalescence event has occurred, extending Δ*t* to the next time on the predefined grid in which a coalescence has occurred. The estimator is also undefined if 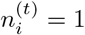 because the expected number of lineages remaining approaches but never reaches 1. In this case, we assume the coalescent time passed in the interval is 2(1 − 1/*n_i_*(0)), which is the expected amount of time to coalesce from *n_i_*(0) lineages to one lineage. Finally, once a subtree coalesces to one lineage, we assume that population size remains constant into the past for the ancestral allele, and for the derived allele, that it remains constant before dropping to zero at in the middle of the branch connecting the derived subtree to the rest of the tree.

The approximate variance in Eq. 49 is undefined if the estimated allele frequency 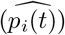 is equal to 0 or 1—in which case we define the estimated variance of 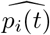 to be 0. It is also undefined if only one lineage remains on either background at the more recent end of the time interval (*n_i_*(0) = 1 or *m_i_*(0)=1). If one lineage remains on one of the backgrounds, then we estimate the variance as the variance of the waiting-time estimator with *ℓ* = 1. If only one lineage remains on each of the two backgrounds, we define the estimated variance of 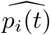 to be 1/12, which is the variance of a Uniform(0, 1) random variable.

### D The correspondence of *T_X_* and *Q_X_*

In this section, we explain the correspondence between the *T_X_* statistic proposed in Eq. 16 and the statistic of Berg and Coop (2014). If the population history is tree-like, then can be thought of as checking for overdispersion— relative to neutral expectations—of polygenic scores at the tips of the population tree. In contrast, *T_X_* checks for overdispersion at a set of timepoints along one branch of a population tree (i.e. a single population at different times).

Briefly, Berg and Coop start with a vector 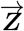 whose elements are population-mean polygenic scores for a set of populations minus an assumed value for the population-mean polygenic score in an ancestral population. Under neutrality, 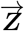 follows a MVN(0, 2*V_A_**F***) distribution, where *V_A_* is the additive genetic variance, and ***F*** is a matrix reflecting the shared history of the populations. In particular, if the population history is tree-like, then entry *F_ij_* in the ***F*** matrix reflects the branch length shared by populations *i* and *j* since their descent from the ancestral population. If 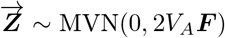, then the transformed vector 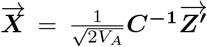—where ***C*** comes from the Cholesky decomposition of ***F***, such that ***F*** = ***CC^T^***—obeys a MVN(0, ***I***) distribution. Thus, the sum of the squared entries of the transformed vector ***X***—the quantity labeled *Q_X_*—obeys a *χ*^2^(*w*) distribution under neutrality, where *w* is the length of 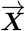.

In the setting of this paper, we have population-mean polygenic scores *Z*(*t*_0_), *Z*(*t*_1_), …, *Z*(*t_w_*) for the population ancestral to one present-day population, assessed at various times. Each *t_j_* represents some amount of time before the present and 0 ≤ *t*_0_ < *t*_1_ < … < *t_w_*. In practice, we set *t*_0_ =0 and treat *Z*(0), the population-mean polygenic score in the present, as Berg and Coop treated the ancestral polygenic score, so

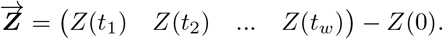

In this setting, with the present-day population serving the role of “ancestor”, the shared branch length for two timepoints along one branch of a population tree is their shared distance backward from the present day, which is to say the branch length from the present back to the more recent of the timepoints.

Thus, for this instance of 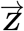, each entry *F_ij_* in the ***F*** matrix is equal to *t*_min(*i,j*)_, meaning

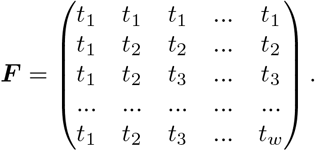

The Cholesky decomposition ***F*** = ***CC^T^*** then gives

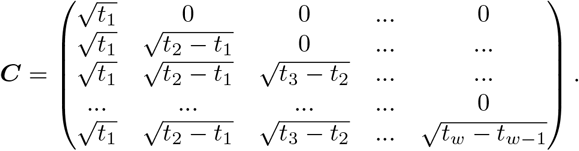

The inverse of this matrix is

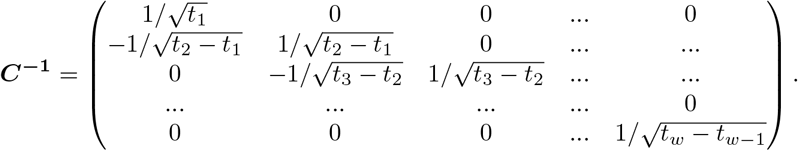

If *V_A_* is a constant, then the entries of 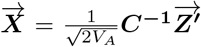 are equal to the *X_j_* values of Eq. 15, and *Q_X_* is equal to the quantity in Eq. 16.

### E Simulation details

We ran simulations of polygenic scores, as well as coalescent trees and haplotypes associated with them, with and without selection.

For each trait, we simulated the frequency of the derived allele at each of *k* independent trait-associated loci. We ensured that at all loci, the minor allele frequency was at least .01, both at the onset of selection (if applicable) and at the present. Allele-frequency time courses were simulated using the normal approximation to the diffusion.

In particular, for each locus, an effect size for the derived allele was drawn from the normal distribution with expectation zero and variance equal to 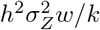 where *h*^2^ is the desired heritability (because we are focused on polygenic scores rather than traits themselves, *h*^2^ = 1), 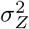 is the desired variance of the trait (i.e. polygenic score)—always 1 in our simulations, *w* is a modified version Watter-son’s constant for the population 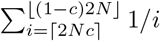 with *c* equal to the minimum minor allele frequency being drawn (here, *c* = .01), and *k* is the number of independent loci affecting the trait. Drawing effect sizes from this distribution gives the property that under neutrality, if the loci affecting the trait are independent, and a trait is formed by adding the individual’s polygenic score to an independent “environmental” term with variance 1 − *h*^2^, then the resulting trait has variance 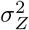 in expectation.

Once the effect size was specified, we selected a derived allele frequency from the population-level neutral site frequency spectrum, conditional on the requirement that the minor allele frequency be at least .01. Under neutrality, this randomly drawn allele frequency was treated as the derived allele frequency at the present. In simulations including a period of selection before the present, this frequency was used as the allele frequency at the start of selection. In simulations with selection, the derived allele frequency was simulated forward in time in steps of 1/(2*N*) coalescent units (i.e. one diploid generation). Conditional on frequency *p_t_*, the frequency at the next time step, *p*_*t*+1_, was drawn from a normal distribution with expectation *p_t_* + *sp_t_*(1 − *p_t_*) and variance *p_t_*(1 − *p_t_*)/(2*N*). Here, *s* is the selection coefficient on the derived allele at time *t*, which is computed as *s* = *αβ* (Charlesworth and Charlesworth, 2010, Eq. 3.17), where *α* is the selection gradient on the trait at time *t* and *β* is the effect size of the derived allele. Two clarifications about the selection coefficient: first, this is the selection coefficient for the heterozygote, meaning that if the fitness of the ancestral homozygote is proportional to 1 (marginalizing over the other loci that affect the trait), then the fitness of the heterozygote is 1 + *s* and the fitness of the derived homozygote is 1 + 2*s*. Similarly, the effect size *α* is the expected trait value difference between the heterozygote and the ancestral homozygote, holding the genotype at all other loci constant. Second, the selection gradient is equal to 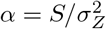, where *S* is the selection differential on the trait—that is, the mean difference in phenotype between the parents of the next generation and the overall population (Charlesworth and Charlesworth, 2010, sec. 3.3.iii)—and 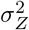 is the trait variance. Here, we used the expected trait variance at the onset of selection (which was 1) to set the value of *α*, which was retained over the course of the period of selection, regardless of the realized value of the trait variance.

Once the allele-frequency time course was simulated forward to the present, it was retained if the minor allele frequency in the present was at least .01. (Otherwise, the time course was discarded and the process was repeated.) To simulate the period from the origin of the mutation up until the onset of selection, we simulated steps backward in time using the normal approximation to the neutral diffusion conditional on loss of the allele. In particular, given a derived allele frequency *p_t_*, the allele frequency 1/(2*N*) coalescent time units further in the past was drawn from a normal distribution with expectation p_t_(1 − 1/(2*N*)) and variance *p_t_*(1 − *p_t_*)/(2*N*) (Przeworski et al., 2005; Berg and Coop, 2015; Lee and Coop, 2017).

In each set of simulations with selection, we retained polygenic-score time courses if their difference in value between the beginning and end of the period of selection was within 5% of the target value, which is given by 2*NδtS*, where *N* is the effective population size, *δt* is the duration of selection in coalescent units, and *S* is the selection differential.

To simulate coalescent trees and haplotypes at each locus, these allele-frequency time courses were used as input to mssel (Berg and Coop, 2015), a version of ms (Hudson, 2002) that can incorporate by conditioning on user-specified allele-frequency time courses. In mssel, we set the sample size to 200 and the number of derived chromosomes to the value of a Binomial(200, *p*) random draw, where *p* is the present-day frequency of the derived allele. If the random draw was equal to 0 or 200, then it was set to 1 or 199, respectively, in order to force the locus to be segregating in the sample. We chose a population size of 10,000, a haplotype length of 200,000 (with the effect locus at position 100,000), a per-base-pair mutation rate of 2 × 10^-8^, and a per-base-pair recombination rate of 2.5 × 10^-8^. These values imply population-scaled ms inputs of -r 199.5 and -t 159.68. Here, 4*NrL* = 200 and 4*NμL* = 160; our slight differences from these values arise from the fact that we compute 4*Nf*(0) where f is the probability distribution function of either a Binomial(*L, r*) random variable or a Binomial(*L, μ*) random variable, respectively, with *L* = 200,000. That is, we define the locus-wide recombination (or mutation) rate as one minus the probability that no recombinations (or no mutations) occur anywhere in the locus. We used the flags -T and -L to include the true trees and times to the most recent common ancestor in the output.

The coalescent trees produced by mssel for the selected sites were used as the true trees, and the haplotypes produced were used as input to RENT+ to generate estimated trees (Mirzaei and Wu, 2016). Both true and estimated trees were read into R (R Core Team, 2013) and handled using the ape package (Paradis et al., 2004). All the estimators and tests used in this paper were coded in R.

### F Power comparison with an analogue of tSDS

Field et al. (2016) proposed the singleton density score (SDS) as a test statistic for detecting very recent selection. Because recent selection distorts the terminal branches of the coalescent at a selected locus, Field et al. reasoned that recent selection would alter the distribution of singleton mutations around selected sites. Specifically, favored alleles are expected to have short terminal coalescent branches and thus relatively few singletons nearby compared with disfavored alleles.

Field et al. use distances to the nearest singletons on each allele’s background to estimate the difference in the mean length of the terminal branches for the two alleles. This estimated mean difference is then standardized by an empirical mean and standard deviation computed from neutral simulations at a similar derived allele frequency. To test for selection on polygenic traits, they construct a trait SDS (or tSDS) by summing each locus’s standardized SDS, signed so that positive values suggest selection for higher trait values.

Here, we compute power for an analogue of tSDS in the simulations shown in Figure 6, with the differences that in our case, the true terminal branch lengths are known, and we use the true effect sizes in computing tSDS. Both of these changes should enhance the power of tSDS. Specifically, at each locus the SDS value is

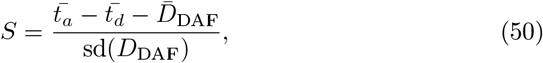

where 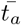 is the mean terminal branch length for the ancestral allele, 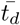 is the mean terminal branch length for the derived allele, 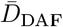 is the sample mean of 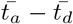 computed from 500 neutral simulations at the same sample size and derived allele frequency (in bins of .005), and sd(*D*_DAF_) is the sample standard deviation of 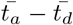 from the same 500 neutral simulations. (It is also possible to replace 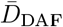 and sd(*D*_DAF_) with analytical values from Fu and Li (1993) under the assumption that the allele frequency does not change. Doing so gives similar results to those we report here.) To compute the tSDS statistic for a polygenic score, we use

**Figure F.8:**
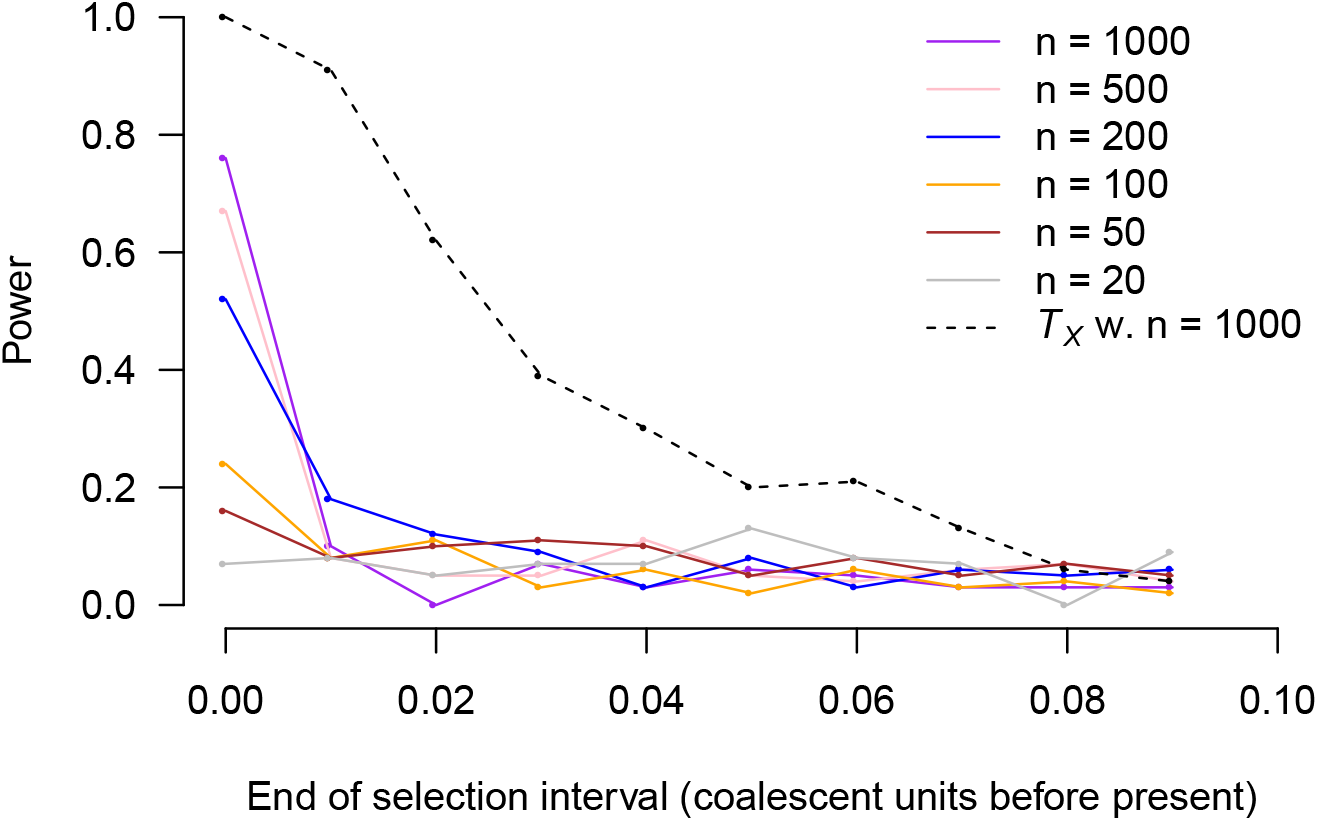
Power of the tSDS analogue as a function of timing of selection and present-day sample size. In addition to results for the tSDS analogue, we show power from the *T_X_* test with a sample of 1000 chromosomes as a guide to the eye for comparisons with Figure 6 (dashed black line). As in Figure 6, power is shown on the vertical axis, and the horizontal axis shows the timing of a pulse of selection lasting .005 units and resulting in an approximate one-standard-deviation shift in the population-average polygenic score (in present-day standard deviations). Each point is based on power from 100 simulations.

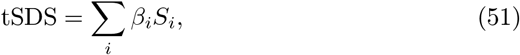

where *β_i_* is the derived-allele effect size at locus *i* and *S_i_* is the value of Eq. 50 for locus i. (In Field and colleagues’ version, *β_i_* is replaced with sign(*β_i_*), which has lower power when the effect size is known.) A polygenic score will tend to have higher tSDS if higher polygenic score values have been selected for recently. To test for significance, we form a permutation distribution by recomputing tSDS 10, 000 times while randomly permuting the effect sizes.

The power of the tSDS analogue as a function of the timing of a selective pulse leading to a ~ 1-standard-deviation change in the polygenic score is shown in Figure F.8. We also show the power of *T_X_* with a sample size of 1,000 chromosomes for comparison (from Figure 6). Compared with *T_X_*, the tSDS has lower power, and its power drops off more rapidly as the timing of selection moves further into the past. This is not surprising because tSDS relies only on terminal branches, and *T_X_* uses all the intercoalescent times.

At this writing, tSDS has the advantage of scaling readily to large samples. In contrast, *T_X_* requires a reconstructed tree, which is currently error prone and computationally intensive. However, as tree reconstruction becomes faster and more accurate, the ability to use information from non-terminal branches will provide power benefits, especially beyond the very recent past.

### G Height analysis details

We estimated the time courses of two of the polygenic scores for height studied in Berg et al. (2018). Information about the loci included in each polygenic score—including rsID, effect size, chromosome, and position—was provided by Jeremy Berg. Each polygenic score was initially constructed by choosing the locus with the lowest p value for a test of association with height—conditional on a minor allele frequency of at least 5%—within each of ~ 1700 approximately independent genomic regions defined by Berisa and Pickrell (2016). (For GIANT, the polygenic score includes 1697 loci, and for UK Biobank, 1700 loci are included.) For the GIANT polygenic score, effect sizes and *p* values were taken from Wood et al. (2014), and for the UK Biobank polygenic score, effect sizes and *p* values were taken from Churchhouse et al. (2017). (Berg et al. (2018) includes several polygenic scores constructed from UK Biobank effect sizes; the one we use corresponds to their “UKB-GB.”)

We used sequence information from the 1000 Genomes GBR subsample (1000 Genomes Project Consortium, 2012, release 20130502), which includes 91 genomes. For each locus included in each polygenic score, we extracted phased sequence information in a window extending 100,000 bases from the locus on each side using tabix (Li, 2011). The resulting .vcf file was processed into a form acceptable by RENT+ using vcftools (Danecek et al., 2011) and R (R Core Team, 2013). We then used RENT+ (Mirzaei and Wu, 2016) to estimate an ARG for the region, including the “-t” flag to estimate branch lengths. Our version of RENT+ was modified slightly to print branch lengths to greater precision and to print RENT+’s internal estimate of *θ* that it uses to scale the branch lengths.

For both polygenic scores, we rescaled the effect sizes so that their standard deviation in the present-day sample is 1, assuming linkage equilibrium among loci. In particular, if the sample frequencies of the effect alleles at the *k* loci are *p*_1_, *p*_2_, …, *p_k_* and the effect sizes are *β*_1_, *β*_2_, …, *β_k_*, then if the loci are independent, the variance of the polygenic score *Z* is

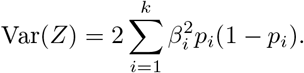

If Var(*Z*) is computed from the original effect sizes, then we rescale the effect sizes as 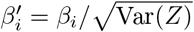 when computing the polygenic scores.

Finally, we rescaled the branch lengths estimated by RENT+. RENT+ estimates a per-nucleotide population-scaled mutation rate *θ* = 4*Nμ* using Watter-son’s estimator (Watterson, 1975). That is, for a sample of n haplotypes each covering a region of w base pairs, with S the number of segregating sites in the sample, *θ* is estimated as

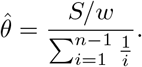

The time in coalescent units separating a pair of haplotypes is then estimated as 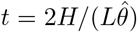, where *H* is the Hamming distance between a pair of haplotypes. For samples of size greater than two, RENT+ estimates branch lengths using a similar logic but using local distance matrices that take into account inferred recombinations.

We scaled RENT+’s estimated times because the Watterson estimator is biased downward if the population has been growing exponentially, and the human population has grown super-exponentially (Keinan and Clark, 2012). As a result, the inferred coalescent times from RENT+ were implausibly large. We first multiplied branch lengths at each locus by the *θ* estimate at each locus, giving branch lengths in units of twice the mutational distance per base pair at all sites. To convert these mutational distances into approximate coalescent units, we computed the time to the most recent common ancestor (tMRCA) at each locus, and then rescaled the branch lengths at all loci by a constant factor that set the mean tMRCA to 2, the expectation in coalescent units under neutrality.

